# Lab-on-a-chip device integrating a human stem cell-based blood-brain barrier model with neural organoids for translational research

**DOI:** 10.64898/2026.09.15.750866

**Authors:** Judit P. Vigh, Anna E. Kocsis, Ana R. Santa-Maria, Nóra Kucsápszky, Emese Bató, Silvia Bolognin, Jens C. Schwamborn, András Kincses, Sándor Valkai, Anikó Szecskó, Szilvia Veszelka, Mária Mészáros, Gergő Porkoláb, Thi Ha My Phan, Jeng-Shiung Jan, Roland Wirth, Zoltán Szabó, Maxime Culot, Yoichi Morofuji, András Dér, Mária A. Deli, Fruzsina R. Walter

## Abstract

For advanced biomedical research on central nervous system pathologies and brain targeting of nanotherapeutics, complex and well described human blood-brain barrier (BBB) models are crucial. Here we established and characterized a complex lab-on-a-chip (LOC) system integrating a BBB model with neural organoids (NO). We optimized the conditions for BBB-NO models under static and dynamic conditions. The combination of a human stem cell-derived BBB co-culture model with human midbrain organoids in a microfluidic LOC allowed the observation of BBB and neural tissue changes and the separate analysis of the barrier and brain units. The LOC design enabled phase contrast and fluorescent microscopy on the whole brain endothelial culture surface, barrier integrity and permeability measurements across the BBB model, and molecule passage into NOs. BBB-specific endothelial morphology, gene and protein expression, and good barrier integrity were demonstrated, corroborating the strength of the LOC engineering and cellular design. The transport of targeted nanoparticles across the BBB model followed by their entry to neural organoids validated the barrier integrity and transporter functionality of the dynamic, integrated and complex model. As a proof-of-concept experiment to confirm the translational value of the BBB-NO model integrated in the LOC device, we examined a clinically used hyperosmolar iodinated contrast agent, iopamidol, with confirmed neurological side effects. We corroborated that iopamidol induces transient BBB dysfunction and neural effects in the complex system, not only validating the complex dynamic BBB-NO model but also pointing to the necessity to study BBB changes together with NO functions.

## Introduction

Studying the role of the blood-brain barrier (BBB) in central nervous system (CNS) pathologies and brain targeting of nanotherapeutics are important areas in the field of biomedicine [1,2]. Balanced function of cerebral microvascular endothelial cells (EC), forming the anatomical basis of the BBB, is crucial to keep brain homeostasis [3]. Disturbance in cell-cell interactions in the neurovascular unit, EC junctional connections and EC transporter activity all lead to or are a consequence of neurological deficits in a large number of CNS pathologies [4]. Specific transporters of brain ECs, low number of intracellular vesicles, functional glycocalyx and good barrier properties are all key features of an *in vitro* BBB model used for drug delivery and CNS disease studies. Therefore, the establishment of high fidelity, complex, reliable and reproducible, preferably human cell type-based *in vitro* BBB models have never been timelier and more critical [5,6].

Cultured brain ECs were first developed in the late 1970s and early 1980s [7,8]. Since then, primary models of rodent, bovine or porcine origin and brain EC cell lines were frequently used as BBB cell culture systems, while in the past ten years human hematopoietic and induced pluripotent stem cell (iPSC)-derived brain endothelial cells are in the focus of investigations and modelling [6,9]. Static *in vitro* BBB models created on cell culture inserts, enabling co-culture of multiple cell types and cell-cell communication through secreted factors, are the most used systems to study BBB physiology, pharmacology and pathology [9,10]. In the past decade, to mimic physiological blood flow, spotlight turned on lab-on-a-chip (LOC) devices that allow dynamic modelling by incorporating fluid flow-induced shear stress to the surface of culture ECs in a 3D setting [5,11]. Complex microphysiological systems were established in chip devices with dual- or multi-compartments and built-in biosensors that allow the culture of brain ECs with other CNS cell types like astrocytes, brain pericytes, microglia and neurons [5,10].

Human stem cell-based organoids provide a versatile tool to model complex diseases, cell-cell communication, development and drug targeting *in vitro* facilitating the translation from the laboratory results to the clinical setting [12]. Parallel with the advancements in the iPSC and microphysiological system fields, the first cortical brain organoids were also differentiated more than ten years ago [13]. Although neural organoids (NO) with limited vasculature exist, currently no models are available with perfusable vessels and mature BBB properties [14].

For advanced biomedical applications with the goal of studying cerebral vessel-related complications in CNS diseases and drug or nanomedicine delivery, new cell culture-based BBB models with a higher complexity are needed. Modelling the BBB with multicellular *in vitro* assays, investigating pathologies at the brain EC level and studying drug transfer across the barrier is only the first step. The question to what happens subsequently in the brain tissue is yet to be investigated with advanced *in vitro* systems, such as neural organoids [6,15]. Dynamic BBB models established together with neural cell types has been published, but in these models the neural compartment (i) lacks structure, (ii) is mixed with other cell types, like astrocytes, pericytes or microglia in a 2D environment, (iii) is missing complex cell-cell interactions such as in neural organoids, or (iv) the model does not possess pump-driven laminar perfusion and/or sensors detecting barrier impedance [16]. Currently no complex LOC system exists that allows the detailed characterization of the BBB specific features, like morphology, barrier integrity, brain EC gene and proteomic expression, as well as biomedical applications simultaneously on the BBB and NO models such as nanoparticle transport and translational research.

In this work we aimed to fill the gap in the landscape of complex LOC systems integrating a BBB and a neural model. Here we present the successful assembly and optimization of a BBB-NO model under static and dynamic circumstances. The dynamic model was based on our previous versatile-LOC system [17,18,19] that was improved to integrate neural organoids. The unique combination of a human BBB co-culture model with human iPSC-derived midbrain organoids in a laminar flow-based two-compartment system allows the in-depth observation of BBB and neural tissue changes and the separate analysis of the barrier and brain units. The BBB-NO-LOC design enables phase contrast and fluorescent microscopy on the whole culture surface, barrier integrity and permeability measurements across the BBB model and molecule passage into neural organoids. Our experiments demonstrated that this model has brain EC specific morphology, gene and protein expression, and good barrier integrity with functional transport. The experimental design also confirmed the translational value of the BBB-NO-LOC not only validating it, but pointing to the necessity to study BBB changes together with neural organoid functions.

## Materials and Methods

### Materials

All the materials used were purchased from Merck Life Science Kft., Budapest, Hungary, unless otherwise indicated.

## Methods

### Lab-on-a-chip design and fabrication

The lab-on-a-chip (LOC) device used here consists of two poly(dimethylsiloxane) (PDMS, Sylgard 184, Dow Corning GmbH, Germany) channels, separated by a porous polyester (PET) membrane (0.45 µm pore size, 2 × 10 pores/cm², and 23 µm thickness; It4ip, Belgium) (Fig. 4B). The biochip was fabricated as described in our previous publications [17,18,19], with some modifications to allow the insertion of NOs into the system. In this device, three holes were drilled into the bottom part of the LOC for the organoid holder caps, and these were glued into the holes using Norland optical adhesive 81 (Norland Products, USA). The electrode structure was also modified. Gold/platinum (Au/Pt) electrodes were designed to perform electrical impedance spectroscopy. First, Au electrodes were sputter-coated for 80 s at 50 mA, followed by the Pt electrodes under the same conditions. Copper wires were glued to the electrodes using a two-component electrically conductive epoxy (CW2400, ITW Chemtronics, USA). The top and bottom PDMS channel sandwich was spin coated with an additional thin layer (20–25 µm) of PDMS to provide additional sealing to prevent medium leakage. The LOC was then assembled and sterilized oxygen plasma treatment for 10 min, and then addition of 70% ethanol for 1 h. After sterilization, the devices were washed five times with sterile distilled water before coating and cell seeding.

### Cell cultures

#### In vitro BBB model

The BBB model consists of human endothelial cells (human brain-like ECs) in co-culture with brain pericytes [20]. The human brain-like ECs were derived from CD34+ cord blood hematopoietic stem cells through a 15–20-day differentiation period using VEGF_165_ [21]. The protocol was approved by the French Ministry of Higher Education and Research (ethics approval no. CODECOH DC2011-1321) in accordance with the World Medical Association Declaration of Helsinki. Informed consent was obtained from the donors’ parents. After differentiation, human brain-like ECs were seeded onto collagen type I-coated culture dishes and maintained in endothelial cell culture medium (ECM, ScienCell, USA) supplemented with 5% fetal bovine serum (FBS, ScienCell), 1% endothelial cell growth supplement (ECGS, ScienCell), and gentamicin (50 µg/ml). After reaching confluency, human brain-like ECs were trypsinized and frozen for later use. In all experiments, human brain-like ECs were used at passage 7. To reach the brain-like endothelial phenotype, human brain-like ECs were co-cultured with brain pericytes [21]. For experiments where human brain-like ECs were grown alone, pericyte conditioned medium was added in a 1:1 ratio.

Bovine brain pericytes were seeded to collagen type I-coated culture dishes and maintained in Dulbecco’s modified Eagle’s medium (DMEM, low glucose, Gibco, Thermo Fisher Scientific, Waltham, MA, USA) supplemented with 20% fetal bovine serum (FBS), 1% Glutamax (Life Technologies, Carlsbad, CA, USA), and gentamicin (50 µg/ml). The confluent cell layer was then trypsinized and seeded onto the experimental surface at passage 12–13. Cells were kept in a humidified incubator at 37 °C with 5% CO_2_.

#### Neural organoids

The generation and maintenance of the neural organoids were described by Nickels et al (2020). Briefly, neural organoids were differentiated from human floor plate neuronal progenitor cells derived from a healthy donor and mainly contain midbrain-specific dopaminergic neurons [22]. All work with human stem cells was done after approval of the national ethics board, Comité National d’Ethique de Recherche (CNER), under the approval numbers 201305/04 and 201901/01. neural organoids reached maturity after 30 days of differentiation (DoD) and were kept in ultra-low attachment 96-well plates (Corning, New York, NY, USA) in a 1:1 mixture of Dulbecco’s Modified Eagle Medium/Nutrient Mixture F-12 (DMEM/F12, Gibco) and Neurobasal medium (Gibco). This was supplemented with gentamicin (50 µg/ml), L-glutamine (2 mM, Thermo Fisher Scientific), B27 supplement (100×, Gibco), N2 supplement (200×, Gibco), dibutyryl cAMP (500 µM), brain-derived neurotrophic factor (BDNF, 10 ng/ml, Peprotech, Thermo Fisher Scientific), glial cell-derived neurotrophic factor (GDNF, 10 ng/ml, Peprotech), transforming growth factor-β3 (TGF-β3, 1 ng/ml, Peprotech), Activin A (2.5 ng/ml, Peprotech), L-ascorbic acid (200 µM), and γ-secretase inhibitor (DAPT, 10 µM, Tocris Bioscience, Bristol, UK). The maturation medium was changed three times per week, and neural organoids were used for experiments between DoD 55 to 120.

#### Static cell culture model

First, cells for the cell culture insert-based static BBB model were seeded: brain pericytes were added at a concentration of 5 × 10³ cells on the bottom of collagen type IV (100 µg/ml)-coated polycarbonate cell culture inserts (PC, 0.4 µm pore size, Cat. No. 3413, Costar). After the adhesion of the pericytes, inserts were turned into 24-well plates (Corning Costar) and human brain-like ECs were seeded to the top of the cell culture inserts coated with collagen type IV (100 µg/ml) and fibronectin (25 µg/ml), at a density of 1.3 × 10^4^ cells. During the 7 days of co-culture, cells were kept in a CO_2_ incubator at 37 °C with 5% CO_2_ (Fig. 1).

**Fig. 1.**
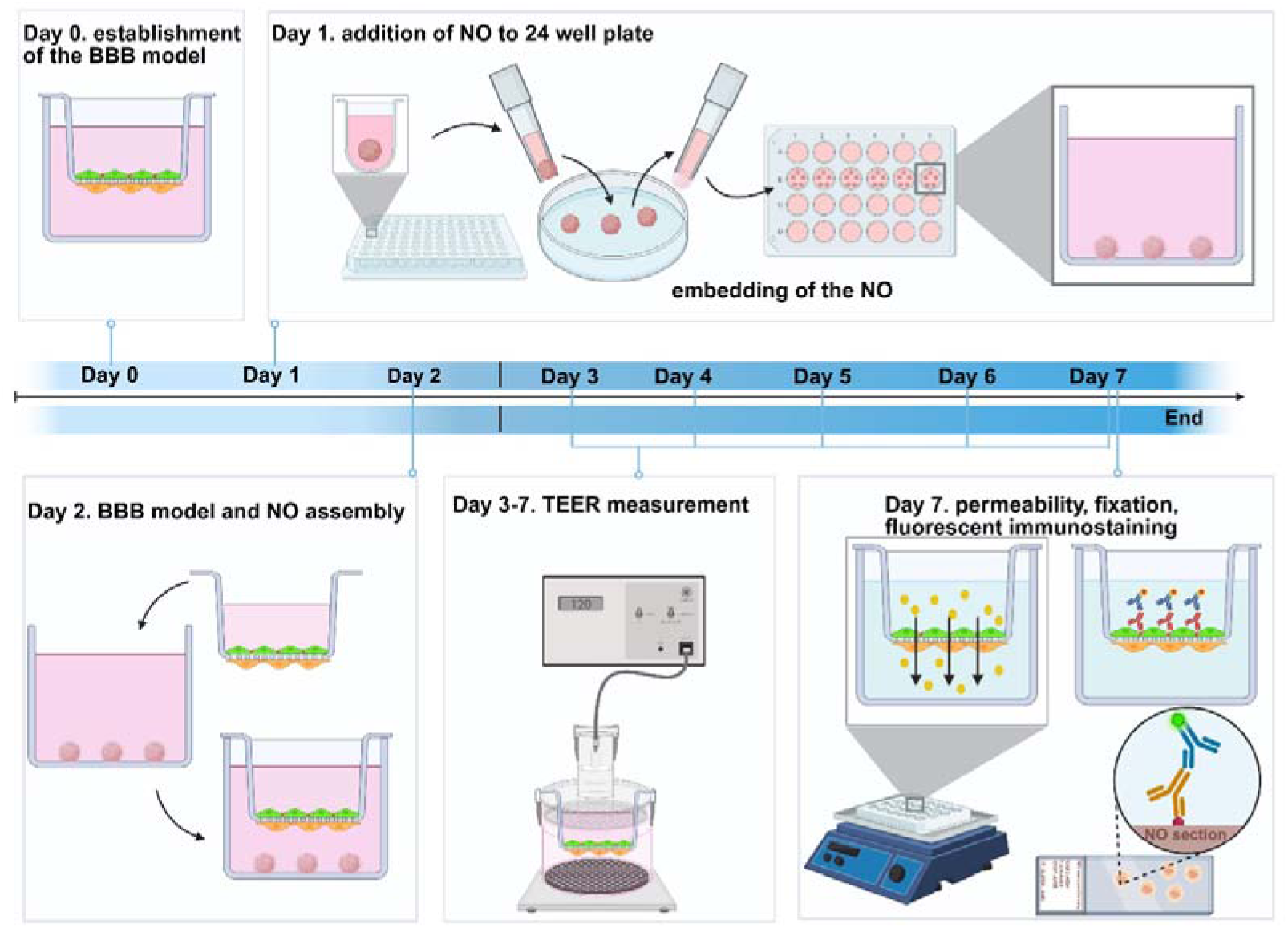
Timeline of the static BBB-neural organoid (BBB-NO) co-culture model establishment. On day 0, the BBB model was established by co-culturing human brain-like ECs (green) and brain pericytes (orange) on opposite sides of a cell culture insert. On day 1, neural organoids (NO) were transferred and embedded in growth factor-reduced Matrigel at the bottom of a 24-well plate. On day 2, the BBB-NO model was assembled by placing the cell culture insert into the well containing the embedded organoid, with a 1:1 mixture of BBB cell culture medium and maturation medium added to the bottom compartment. From day 3 to day 7, trans-endothelial electrical resistance (TEER) was measured daily to monitor barrier integrity. On day 7, permeability assays, cell fixation, and fluorescent immunocytochemistry were performed on the assembled model, including immunostaining of NO sections. Created in BioRender. Vigh, J. (2027) https://BioRender.com/zo00mt2

Neural organoids were embedded in growth factor-reduced Matrigel (Corning) at the bottom of a 24-well plate (Corning Costar), as described previously [23]. The BBB-NO model was assembled on the second day of the co-culture of the BBB model, by adding a 1:1 mixture of BBB cell culture medium and maturation medium to the bottom compartment. The top compartment received human brain-like ECs cell culture medium. The medium was changed every second day, and the model was kept under these conditions for 5 more days (total of 7 days). Trans-endothelial electrical resistance (TEER) was measured every day. On Day 7 permeability measurements for fluorescent marker molecules were performed. Cells were fixed and stained with immunocytochemistry for BBB specific markers (Fig. 1).

#### Dynamic cell culture model

Human brain-like ECs were co-cultured with brain pericytes in the LOC system similarly to the static cell culture model. Here, 6 × 10 human brain-like ECs and 2.5 × 10 brain pericytes were seeded onto the two sides of the polyester membrane of the biochip (Fig. 2), which was coated with collagen type IV (100 µg/ml) and fibronectin (25 µg/ml) from the top for human brain-like ECs, and collagen type I (150 µg/ml) from the bottom for pericytes. Until the second day of the co-culture of BBB model, both compartments received endothelial medium. The LOC device with the cells was kept at 37 °C in a humidified incubator with 5% CO_2_.

**Fig. 2.**
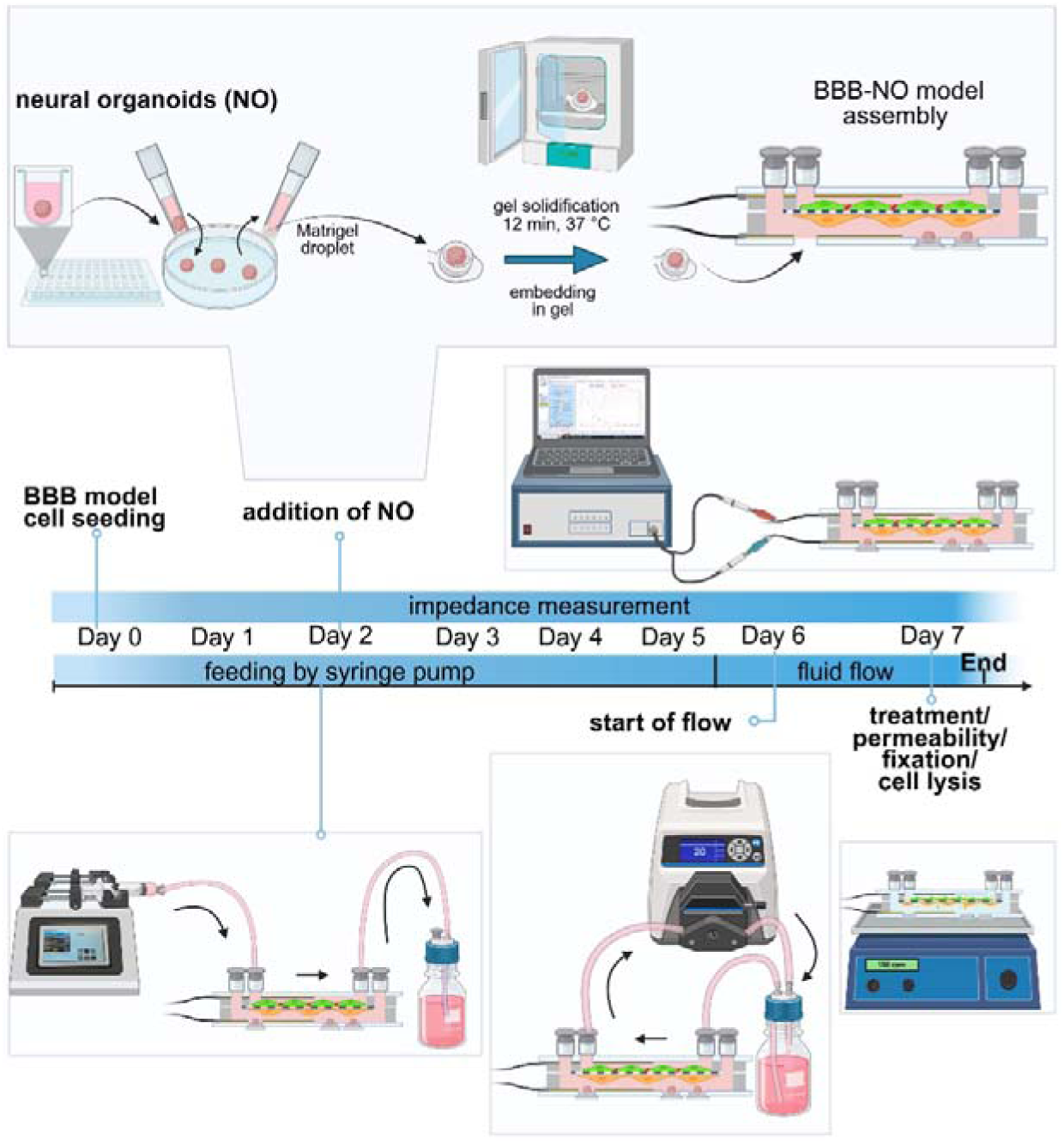
Timeline of the dynamic BBB-neural organoid (BBB-NO) lab-on-a-chip (LOC) model establishment. On day 0, the BBB model was assembled by co-culturing human brain-like ECs and brain pericytes on opposite sides of the polyester membrane within the LOC device. Neural organoids (NO) were embedded into holder caps in a Matrigel droplet, followed by gel solidification for 12 min at 37 °C, and were then incorporated into the LOC device on day 2, completing BBB-NO model assembly. From day 0, the upper channel was fed automatically by a syringe pump, and impedance measurements were performed daily. On day 6, dynamic flow conditions 1 ml/min (0.4 dyne/cm²) were initiated using a peristaltic pump. On day 7, the LOC devices were used for functional tests, permeability assays, cell fixation, or cell lysis for further analyses, depending on the experimental endpoint. Created in BioRender. Vigh, J. (2026) https://BioRender.com/zo00mt2

The neural organoids were embedded in their holder caps with Matrigel/Geltrex (Gibco) on the second day of the BBB model, and from that point, a 1:1 mixture of BBB cell culture medium and organoid maturation medium was used in the bottom compartment. Medium supplementation was performed automatically by a syringe pump (Legato 110, KD Scientific, Holliston, MA, USA) in the upper channel three times per day at a 500 µl/min feeding rate, while in the bottom channel medium was changed once per day manually. On the sixth day of the BBB-NO model co-culture, dynamic conditions were applied using a peristaltic pump (Masterflex, Cole-Parmer, USA) for 24 hours at a flow rate of 1 ml/min (0.4 dyne/cm²) (Fig. 2). Every day of the co-culture, impedance of the human brain-like ECs layer was registered. On Day 7 experiments were performed, permeability of fluorescent marker molecules was measured, and cells were fixed for subsequent analysis.

#### Western blot on neural organoids

The effect of human brain-like ECs culture medium was tested on neural organoids, mixed in different ratios with neural organoid maturation medium. The experiments were performed on DoD 20 and 29, since neural organoids are usually ready to use in experiments from DOD 30 (Nickels et al, 2020). Organoids at DoD 20 and 29 were rinsed once with 1x PBS, then neural organoids were lysed in cytoplasmic lysis buffer containing 2% Triton X-100 and complete protease inhibitor (1:25) and phosphatase inhibitor cocktail (1:50) in phosphate buffer (PBS) (Thermo Fisher Scientific). Organoids were introduced into tubes with ceramic beads (Precellys Keramik Kit) and tissueLyser (Quiagen, Germany) was used (5min, 50 oscillation). The lysate was centrifuged for 30 min, 13,000 rpm at 4°C, then the supernatant was transferred to a new tube. BCA assay (Thermo Fisher Scientific) was used according to the manufacturer’s instructions to determine the protein concentration in each sample. Lysates were transferred from the gel to polyvinylidene fluoride membranes in an iBlot2 device (Thermo Fisher Scientific). Membranes were blocked for 1 h at room temperature (RT). Primary antibodies tyrosine hydroxylase and GAPDH were incubated at 4°C overnight. Secondary antibodies were incubated for 1 h, also at RT. Membranes were revealed using the SuperSignal West Pico Chemiluminescent Substrate (Thermo Fisher Scientific).

### Barrier integrity tests

#### Trans-endothelial electrical resistance

Trans-endothelial electrical resistance of the static BBB-NO co-culture was measured daily using a chamber electrode connected to a 4-channel voltohmmeter (EVOM2, World Precision Instruments, USA). A heating pad was used during the measurements to sustain the temperature of the system to avoid fluctuations during the measurement [24]. The resistance of inserts without cells was measured as background resistance, and the surface area of the insert was used to express the TEER value (Ohm × cm²).

#### Electrical impedance spectroscopy on the LOC device

Electrical impedance spectroscopy (EIS) measurement was performed in the LOC system every day, before the daily change of culture medium. The LOC devices were removed from the incubator and were placed onto a heating pad set to 37°C for the impedance measurements, ensuring that the measurements were always performed at the same temperature. Since EIS scans span a wide frequency range, from low (mHz) to high (MHz), most processes—from the slowest to the fastest—can be characterized within a single scan. A potentiostat (SP-150, BioLogic, France) was used with the following measurement parameters: the signal was recorded from 10 Hz to 1 MHz, and the amplitude of the sinusoidal excitation signal was 10 mA. The average of three measurements per frequency was taken as the basis for the entire spectrum, with sampling performed six times per decade.

The measurement results were processed using EC-Lab® (BioLogic), a software package for planning, performing, and graphically displaying electrochemical measurements. The resistance of the monolayer was determined according to the following formula:

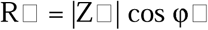

where R□is the resistance of the monolayer (ohm), |Z□| is the magnitude of the complex impedance (Ohm), and φ□ is the phase angle of the complex impedance (rad).

On the microfluidic chip devices, impedance (Z□/Ohm) was measured after collagen coating but before cell seeding, serving as a background value. The impedance values of the culture (Z□/Ohm) were then recorded daily, and the corresponding background values were subtracted to obtain the so-called cell index (CI/Ohm), according to the following formula:

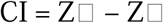

Then the CI was multiplied by the surface of the cell culture membrane to get the impedance (Ohm × cm^2^).

#### Permeability

To assess the barrier integrity of the static and dynamic BBB-NO models, permeability assays for fluorescein isothiocyanate-labeled dextran (MW: 4.4 kDa, FD4) and Evans blue-labeled albumin (MW: 67 kDa, EBA) were performed as described in our previous work [17]. The marker molecules were diluted in Ringer-Hepes solution supplemented with 1% Insulin-Transferrin-Selenium (ITS, insulin 5 µg/ml, transferrin 5 µg/ml, sodium selenite 5 ng/ml; Gibco) and 1% bovine serum albumin (BSA, Thermo Fisher Scientific). The final concentrations were 100 µg/ml for FD4 and 170 µg/ml for EBA.

The cell culture inserts within the 24-well plates and LOC devices were incubated during the permeability assays on a horizontal shaker (150 rpm; Biosan, Latvia) in a CO_2_ incubator at 37 °C for 15 min. FD4 and EBA concentrations in the lower compartment were determined using a Fluorolog 3 spectrofluorometer (Horiba Jobin Yvon; FD4: ex. 485 nm, em. 520 nm; EBA: ex. 582 nm, em. 680 nm). For both the insert-based models and the LOC devices, apparent permeability coefficients (P_app_) were calculated as described previously [19].

#### Real-time cell impedance measurement in a multiwell plate

To measure the growth of human brain-like ECs and the formation of the barrier in the presence of different percentages of neural organoid maturation medium with human brain-like ECs culture medium, we performed a real-time, non-invasive impedance-based measurement (xCELLigence RTCA SP, Agilent, Santa Clara, CA, USA; [25]). For this assay human brain-like ECs were cultured in 96-well plates equipped with integrated gold electrodes (E-plate 96; ACEA Biosciences, USA). The instrument continuously records impedance values at a frequency of 10 kHz, every 10 minutes, and automatically calculates the cell index for each measurement time point based on the equation (Rn-Rb)/10. Rn is the impedance measured between the cells and the electrode, while Rb is the background impedance of the cell-free culture medium [26]. The cell index was normalized to the last time point before treatment in each experiment. E-plates were coated with a mixture of collagen type IV (100 μg/ml) and fibronectin (25 μg/ml), followed by 20 minutes of UV treatment, after which the plate was filled with culture medium and the background impedance was measured. Endothelial cells were seeded to the plates at a density of 4.5 × 10³ cells/well. The culture medium was changed every two days until the cells had grown to confluence and reached the plateau phase. We examined the cells’ response to the organoid maturation medium in two experimental setups. First, upon reaching the plateau phase (Fig. 3A), the endothelial cells were treated with 100-75-50-25% neural organoid maturation medium mixed with endothelial cell culture medium. Triton X-100 solution (1%) served as a control for full cell death. Cell response was monitored for 48 hours. In another experimental setup, cells were treated with a similar timeline to the BBB-NO LOC assembly: cell were treated on the second day after seeding to determine whether the presence of the neural organoid medium affects the growth of endothelial cells (Fig. S1C).

**Fig. 3.**
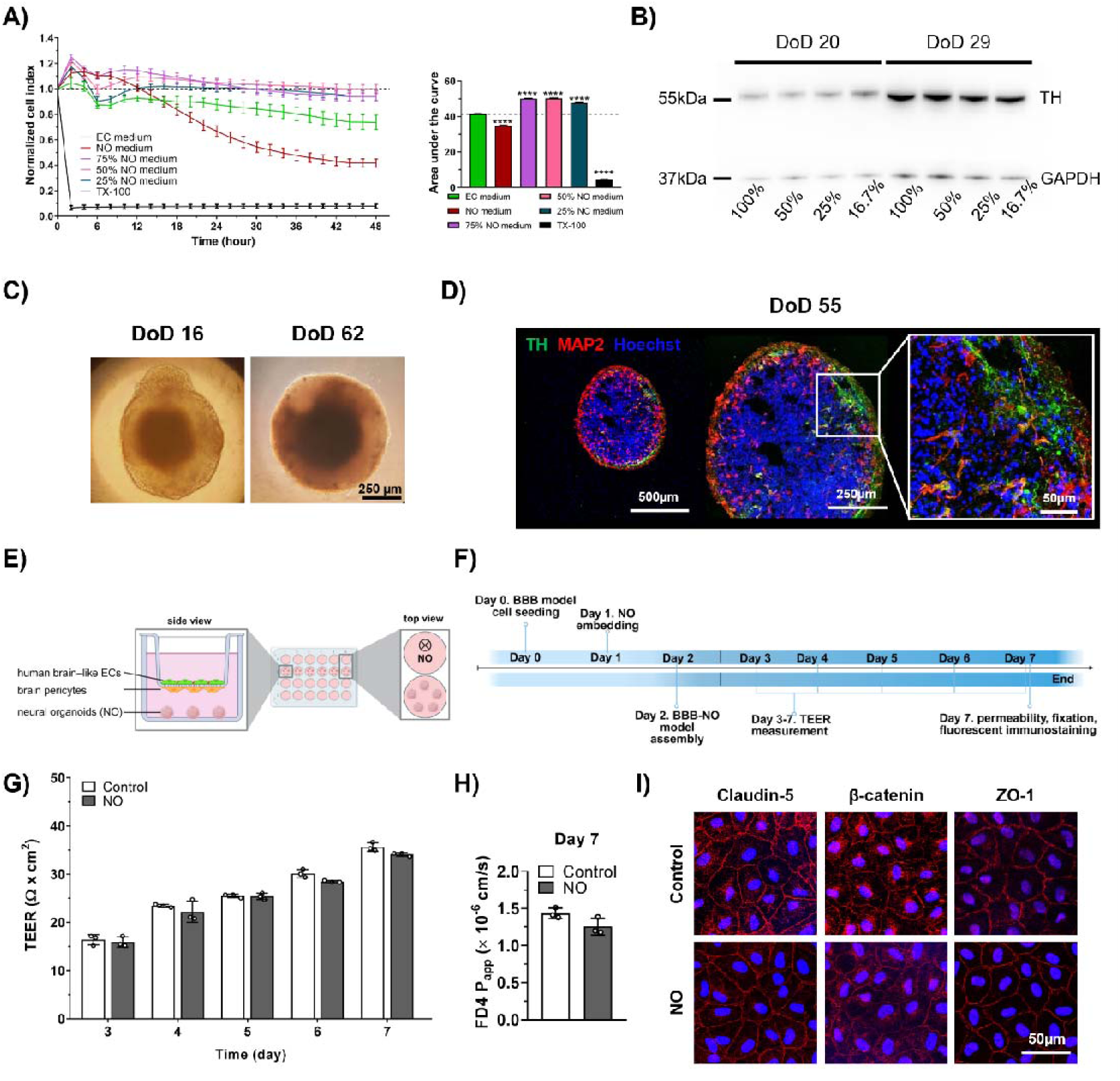
Establishment of the static blood-brain barrier-neural organoid (BBB-NO) model. (A) Real-time impedance analysis on human brain-like ECs of different percentage (75%, 50%, 25%) of NO medium mixed with EC medium. Normalized cell index and area under the curve. TX-100: Triton X-100. One-way ANOVA, Bonferroni post-test, mean ± SD, **** *P*<0.0001, n = 3-10. (B) Western blot of tyrosine-hydroxylase (TH) performed on lysed NOs at Day 20 and Day 29 of differentiation (DoD). NOs were treated for 7 days with different NO medium concentrations (75%, 50%, 25%) mixed with hEC medium. (C) Phase-contrast image of NOs at different maturation stages. 16 days-old and 62 days-old NOs, scale bar: 250 µm. (D) Representative fluorescent microscopic images of 55 days-old NOs. Green: TH, red: microtubule-associated protein 2 (MAP-2), blue: nuclei (Hoechst 33342). (E) Schematic drawing of the static BBB model with or without NOs. (F) Timeline of the assembly for the static BBB-NO model and subsequent analysis. TEER: transendothelial electrical resistance. (G) TEER of BBB model with NOs between Day 3-7. Mean ± SD, n=3. (H) Fluorescein isothiocyanate 4 kDa (FD4) permeability on the BBB-NO system on Day 7. Mean ± SD, n=3. (I) Immunostaining for tight junction proteins. Red: claudin-5, β-catenin and ZO-1, blue: Hoechst 33342, scale bar = 50 µm.

### Immunocytochemistry

#### BBB model

To examine the morphology of the human brain-like ECs in both the cell culture insert and LOC models, cells were fixed with ice-cold methanol for 2 min and immediately washed with PBS (KCl 2.7 mM, KH_2_PO_4_ 1.5 mM, NaCl 136 mM, Na_2_HPO_4_ × 2 H_2_O 6.5 mM, pH 7.4) containing 1% FBS to rehydrate the membrane, or fixed with 3% formaldehyde solution (PFA, Electron Microscopy Sciences, Hatfield, PA, USA) for 15 min at RT, then washed 3 times with PBS. For F-actin staining Alexa Fluor 488 labeled Phalloidin was used. Non-specific binding sites were blocked with 3% BSA for 1 h at RT. Rabbit anti-claudin-5, rabbit anti-β-catenin, and rabbit anti-ZO-1 primary antibodies (Table S1) were used at 1:300 dilutions overnight at 4 °C. On the second day of immunocytochemistry, cells were washed with PBS 3 times before applying the anti-rabbit secondary antibody at 1:400 and Hoechst 33342 nucleus dye at 1:500 dilution (Table S1) for 1 h at RT. Cells were washed again 3 times with PBS and once with distilled water before mounting with Fluoromount-G mounting solution (Southern Biotech, AL, USA).

#### Neural organoids

To characterize neural organoids in the BBB-NO-LOC device, they were fixed with 3% PFA for 75 min at RT then washed with PBS 3 times. Before cryosectioning, organoids were dehydrated with 30% sucrose solution overnight at 4 °C. Organoids were embedded into OCT (Tissue-Tek, Sakura Finetek, Torrance, CA, USA) on dry ice, then 20 µm thin sections were made by using a cryostat (Leica CM 1860, Leica Microsystems, Germany, Veszelka et al. 2022) and placed onto a charged plus glass slide (SuperFrost, Epredia, Seattle, WA, USA). Sections were washed with PBS to remove the OCT and permeabilized with 0.2% Triton X-100 in PBS for 10 min at 4°C. Sections were blocked with 2% normal horse serum (NHS) and 0.3% BSA for 1 h at RT. Neurons were marked with mouse anti-TUJ1 (1:1000, Thermo Fisher Scientific), chicken anti-microtubule associated protein-2 (MAP2, 1:2000, Abcam, Cambridge, UK), mouse anti-tyrosine hydroxylase antibody, astroglia cells were marked with goat anti-glial fibrillary acidic protein (GFAP, 1:500, Abcam) primary antibody (Table S1) in blocking solution at 4 °C, overnight. The next day the sections were washed 3 times with PBS. Then anti-mouse/rabbit/goat Alexa 488, anti-mouse Alexa 555 and anti-chicken Alexa 594 secondary antibodies and Hoechst 33342 nucleus dye (Table S1) were used for 1 h at RT. Sections were washed again with PBS and distilled water then sections were mounted.

#### Image analyses

Fluorescent images were taken with the Leica TCS SP5 confocal laser scanning microscope (Leica Microsystems, Wetzlar, Germany). For image analyses, the ImageJ software (National Institutes of Health, Bethesda, MD, USA) was used.

#### Total RNA isolation

The BBB-NO co-culture model in the LOC was assembled as written above. human brain-like ECs and brain pericyte contact culture was kept together with the neural organoids for five days. On Day 7 human brain-like ECs were lysed directly with RLT Plus buffer (QIAGEN, catalog no. 1053393), as described in our previous publications (27,28). RNA was isolated using the RNeasy Plus Micro Kit (QIAGEN, catalog no. 74034) according to the manufacturer’s protocol, which contains an integrated genomic DNA eliminator spin column. RNA integrity was analyzed using automated capillary electrophoresis (RNA Pico Sensitivity Assay, LabChip GX II Touch HT instrument, PerkinElmer). Samples were stored at −80 °C until further analysis

#### Library preparation and 3_′_ RNA sequencing

RNA sequencing at GenXPro GmbH, Frankfurt, Germany was performed for genome-wide gene expression profiling. Samples with 1 μg of purified RNA were used for library preparation, where a total of 20 libraries were constructed. Fragmented RNA was reverse transcribed into cDNA, using barcoded oligo(dT) primers containing TrueQuant unique molecular identifiers, followed by template switching. Library amplification was done using polymerase chain reaction (PCR), purified by solid-phase reversible immobilization beads (Agencourt AMPure XP, Beckman Coulter, catalog no. A63882). Sequencing was performed on an Illumina NextSeq 500 platform.

#### Bioinformatic analysis of RNA sequencing data

Approximately 20 million single 75-bp reads were obtained per library. Unprocessed sequencing reads were adapter-trimmed and quality-trimmed using Cutadapt (version 4.6, [29]) with the arguments “-e 0.1 -O 3 -q 20 -m 20 -n 8”. Additional sequencing artifacts were removed from the reads with the arguements “-m 20 -u 4 -a ’A{300}X’ -a ’A{10};o=10’”. FastQC (0.11.9) [30] was used to assess the quality of sequencing reads. Processed sequencing reads were mapped to ENSEMBL_dna of hg38 (Homo sapiens) using Bowtie2 (2.4.4, [31]) with the arguments “--sensitive --local”. Quantification of mapped reads to each gene was performed using HTSeq (version 2.0.2, [32]) with the arguments “-i gene_id -r pos - a 0” and setting strandedness to “no”. MultiQC (version 1.12) was used to create a single report visualizing output from multiple tools across many samples, enabling global trends and biases to be quickly identified. Normalized gene expression values (TPM) were calculated by dividing each count value with the sum of all counts for each sample multiplied by one million. Due to the nature of MACE-seq library preparation, normalization for transcript length is not necessary. Raw counts were geometric-mean-normalized using DESeq2 (version 1.50.2 [33]) Testing for differential gene expression was performed using the DESeq2 Bioconductor package (version 1.50.2 [33]) implemented in R (version 4.5.2). As a result, log2FC [log2(fold change)] and P values were obtained for each gene in the dataset. To account for multiple comparisons, false discovery rate (FDR) was calculated using the Benjamini-Hochberg method. Genes with an FDR < 0.05 and log2FC > 1 or log2FC < −1 were considered to be differentially expressed. To perform functional enrichment analysis, the g:GOSt tool in g:Profiler was used to identify overrepresented Gene Ontology terms.

#### Sample preparation of proteomics

For proteomic analyses human brain-like ECs were grown on LOC devices similarly as before the gene sequencing. On the 7^th^ day of the model human brain-like ECs were lysed directly in the LOC devices by using RIPA buffer completed with PMSF, protease inhibitor cocktail and sodium orthovanadate according to the manufacturer’s instructions (RIPA Lysis Buffer System, Santa Cruz Biotechnology, Germany. Whole-cell lysates were stored at −80 °C until proteomics measurements and further analysis. Protein concentration of cell lysates was determined by the BCA Protein Assay, following the manufacturer’s acetone precipitation procedure at a four-fold dilution. For protein digestion, 10 µg of total protein was subjected to reduction and alkylation (9 mM tris(2-carboxyethyl) phosphine and 40 mM chloroacetamide in TFA) for 5 minutes at 95 °C. Samples were then diluted with water to reach a final concentration of 1 M Tris and 5% TFA. Trypsin was added at a 1:50 enzyme-to-protein ratio, and digestion was carried out overnight at 30 °C. The reaction was terminated by adjusting the mixture to 3% formic acid

#### Proteome Analysis by Liquid Chromatography with Tandem Mass Spectrometry (LC–MS/MS)

Digested peptides were analyzed via LC–MS/MS using a Waters ACQUITY UPLC M-Class system coupled to an Orbitrap Exploris 240 mass spectrometer (Thermo Fisher Scientific, Waltham, MA, USA). Peptides were separated on a C18 analytical column under gradient elution conditions. Data were collected in a data-independent acquisition (DIA) mode, and raw files were processed with DIA-NN 1.9.1 [34] to construct a spectral library and quantify peptide abundances. Precursor identifications were filtered at a 1% false discovery rate (FDR), followed by differential expression analysis with FDR-corrected p-value thresholds. LC–MS data normalization and statistical analysis were performed in R using the MS-DAP package version 1.2.1 [35], applying the ‘vsn’ method for normalization and the built-in ‘deqms’ (1.20.0) and ‘msEmpiRe (0.1.0) algorithm for differential expression with an FDR < 0.05 significance cutoff. Only proteins detected in at least two samples in a sample group with two unique peptides were included in the analysis.

### Proof-of-concept functional tests

#### Nanoparticle penetration experiments

To test physiologically relevant passage of human brain-like ECs-targeted molecules and their entry to neural organoids, two types of nanoparticles (NPs) were used. L-alanine (A) and L-glutathione (GSH)-dual-targeted 3-armed poly(L-glutamic acid) (3-PLG-A-GSH) polymeric nanoparticles were coupled with active agents, such as dopamine (3-PLG-dopa-A-GSH). NP preparation, functionalization, drug coupling, physico-chemical properties, toxicity studies and permeability on a static insert model is detailed in our previous publication [36]. All samples were labeled with N-(2-aminoethyl) rhodamine 6G-amide bis(trifluoroacetate) (R6G). For permeability studies across the BBB-NO LOC model, the system was assembled as described above. 3-PLG-dopa and 3-PLG-dopa-A-GSH nanocarriers were diluted in phenol red-free culture medium (ECM, Sciencell, USA) supplemented with fetal bovine serum (FBS; Sciencell), 1% endothelial growth supplement (Sciencell) at a final concentration of 100 μg/ml and added to the donor compartment. Passage of NPs was tested at a continuous laminar flow (500 µl/ml; total volume: 4 ml). Barrier integrity of the BBB model in the chip was also assessed after the NP permeability for FD4 (100 µg/ml), for 30 min. P_app_ was calculated both R6G fluorescent signal and the FD4 as described above. Organoid uptake of the fluorescent cargo was determined after the lysis of neural organoids with Triton-X-100 (10 mg/ml) in distilled water. Organoids were placed into a small centrifuge tube with 200 µl of lysis solution, and were lysed with the help of a conical pestle. Samples were vortexed, then centrifuged at 13000 g for 5 min (Biofuge Pico, Heraeus, Thermo Fisher Scientific, USA), then BCA kit was used to measure protein content of each organoid. This same lysate was also used to determine the amount of fluorescent R6G in the supernatant by using the Fluorolog 3 spectrofluorometer (Horiba Jobin Yvon; TR: ex. 525 nm, em. 551 nm). The nanogram amount of TR in the system was normalized to microgram protein and targeted and non-targeted samples compared.

Another type of nanocarrier was also tested in the BBB-NO-LOC model to further corroborate the strengths of the model. DSPE-PEG-linked A-GSH targeted niosomes/nanovesicles were prepared similarly as described before by our group [23,37,38,39.]. The passage of targeted nanoparticles (N-A-GSH) and non-targeted nanovesicles (N) with a Texas red-BSA (TR-BSA) cargo were characterized, cellular toxicity optimized and passage across static BBB models measured (Fig. S3). Methods are shared in detail in the Supplementary file. Permeability studies and further analysis was performed as written above for the polymeric system, except that the FD4 permeability was run for 15 min after the 4 h NP passage. TR-BSA amount was determined in the neural organoid lysate by using the Fluorolog 3 spectrofluorometer (TR: ex. 589 nm, em. 609 nm).

#### Experiments with hyperosmolar contrast agent iopamidol on brain endothelial cells and neural organoids

The BBB-NO-LOC model was established as described above (Fig. 2). On day 7, human brain-like ECs were treated with iopamidol, a iodine-containing contrast medium monomer, Oypalomin, Mw: 777.09 g/mol) [40], for 30 min at 50% dilution in human brain-like ECs medium. This contrast medium is a high-osmolar contrast agent, with an osmolality of 812 mOsm/kg H_2_O, which is 2.8-fold higher than that of blood. There were two treatment groups: the “acute” group, in which barrier integrity analysis (FD4 and EBA permeability, and TJ immunostaining) and neural organoid functional tests were performed immediately after the 30 min treatment; and the “recovery” group, in which the treatment medium was replaced with normal cell culture medium, fluid flow was applied for another 24 h, and the same tests were performed. Permeability measurements, fixation of human brain-like ECs and subsequent immunocytochemistry were performed as described above. human brain-like ECs were analyzed with the Leica TCS SP5 confocal laser scanning microscope, and rounded up, detaching cells were identified by H33342 nucleus staining besides junctional identification.

#### Esterase activity assays, calcein-AM and DCFDA

During contrast medium treatment not only barrier integrity was assessed, but as neural organoid functionality tests, esterase activity assays (calcein-AM and DCFDA) were performed. For these measurements, neural organoids were removed from the LOC devices and transferred to low-attachment 96-well U-bottom plates (Corning)., calcein-AM, a cell-permeable green dye (Invitrogen, Thermo Fisher Scientific), was used at a concentration of 2.5 µM, diluted in Ringer-HEPES solution. DCFDA solution (2 µM, Invitrogen, Thermo Fisher Scientific) was supplemented with pluronic acid (200 µg/ml) in Ringer-HEPES solution. The 96-well plate containing the neural organoids was placed into a multiwell plate reader, and green fluorescence was recorded at 485 nm excitation and 520 nm emission. Both assays were measured simultaneously. The measurement was performed over 1 h at a temperature of 37 °C.

#### Statistical analysis

Data were presented as mean ± SD. One-way or two-way ANOVA followed by Bonferroni’s post hoc test were used. Experiments were repeated minimum 2 times except the MACE seq and proteomics on the human brain-like ECs. Outlier analysis was performed if needed by using the Rout or Grubbs tests. Data analysis was performed using GraphPad Prism 8.1 Software (GraphPad Software Inc., La Jolla, CA). Differences were considered statistically significant at P < 0.05.

## Results

### Establishment of the blood-brain barrier and neural organoid (BBB-NO) complex model

#### Optimization of the BBB and neural organoid co-culture system

Neural organoids need differentiation media for their maturation and development [22], while brain ECs and pericytes require culture media with a special composition, different from that of the neural organoids [21]. Since neural organoid and EC characteristics and behavior can be influenced by the different cell culture media components in co-culture, we examined the effect of neural organoid maturation medium (NO medium; 25%, 50%, 75% dilutions in EC medium) on confluent human brain-like EC cultures by impedance-based cell analyses for 48 h. Undiluted (100%) NO medium significantly decreased the impedance of human brain-like ECs without recovery. All three dilutions of NO medium significantly elevated the EC cell impedance (Fig. 3A) indicating a beneficial effect on barrier integrity.

The effects of NO medium diluted in EC medium were also tested on the cellular composition and function of midbrain organoids. The previously characterized neural organoids primarily contain midbrain-specific dopaminergic neurons [22]. Neural organoids were cultured in different concentrations (100%, 50%, 25%, 16,7%) of NO medium, then TH protein content, the marker of dopaminergic neurons, was analyzed by Western blot. Midbrain organoids at day 20 of differentiation produced less TH compared to organoids at day 29 of differentiation. The different dilutions of NO medium did not change the TH level (Fig. 3B), which suggested that midbrain organoid identity was kept when cultured in a mixed medium environment. In subsequent experiments a 1:1 ratio of EC and NO medium was used in the lower compartment for both static cell culture inserts and the LOC device as a co-culture medium. This mixture was additionally tested on the impedance of brain-like ECs during the growth phase and maturation of the barrier. When EC medium was changed to 1:1 mixture of EC and NO medium, impedance of ECs did not differ from the group fed with EC medium only (Fig. S1C), confirming its suitability for the timeline of further experiments.

The diameter of midbrain organoids at day 16 of differentiation varied between 500-750 µm (Fig. 3C). The size of organoids between days 60-120 of differentiation was about 1 mm, and they had a more spherical shape and a denser structure (Fig. 3C). Organoids were characterized by immunostaining for neuronal markers microtubule-associated protein 2 (MAP2) and TH (Fig. 3D). To establish the BBB-NO model, organoids at day 60 to 120 of differentiation were used. Organoids were embedded into growth factor-reduced matrix protein solution in order to keep them in a fixed position during experiments. We observed neural processes growing out of the embedded organoids by phase-contrast microscopy (Fig. S1A) indicating healthy, growing and communicating neurons.

To investigate how the number of organoids in the bottom compartment of the static BBB model affect brain EC barrier functions a 7-day BBB co-culture model was used, as described in our previous publication [19]. A contact co-culture of brain-like ECs with brain pericytes was established, and at day 4 of the BBB co-culture 0, 1, 3, or 5 NOs were placed to the bottom compartments (Fig. S1B). TEER was measured with chamber electrodes to investigate barrier integrity on days 5-7. The number of neural organoids did not have a statistically significant effect on the TEER values of brain ECs, except for the resistance of the group containing 1 neural organoid on day 7, where a decrease was measured compared to the control (Fig. S1B). Since on day 7 of the co-culture the 5 organoids/insert arrangement resembled the control values the most, this setup was used in further experiments.

#### Characterization of the static BBB-NO model

First the human brain-like EC and brain pericyte contact co-culture BBB model was established (day 0) and neural organoids were added to the system on day 2 (Fig. 3F). The BBB-NO co-culture was examined for 5 days, with final assays performed on day 7 of the BBB model (Fig. 3E-F). To study the effect of neural organoids on the barrier integrity of human brain ECs in the complex co-culture model TEER was measured every day. The barrier got tighter steadily over time with or without neural organoids in a comparable way. On day 7 dextran permeability assay was performed. No significant differences were found in TEER and permeability values between the groups cultured with or without neural organoids. Good barrier integrity and low permeability for FD4 was shown in both models indicating that neural organoids did not alter BBB barrier integrity when in static co-culture (Fig. 3G-H). The staining for junctional proteins claudin-5, β-catenin and ZO-1 in human brain-like ECs showed typical morphology and continuous pattern at the cell borders in both groups (Fig. 3I). Based on the TEER and permeability measurements, as well as immunofluorescent staining, human brain-like ECs form a mature, functional barrier in the presence of neural organoids in static condition.

### Characterization of the dynamic BBB-NO model

#### Integration of neural organoids into the dynamic BBB-chip

Despite their versatility, one of the main limitations of cell culture insert-based static BBB models is that they do not provide physiological shear stress at the luminal surface of ECs. In our previous work using a microfluidic LOC device we demonstrated that shear stress promotes the maturation of human brain-like ECs and the development of BBB properties [19]. To create a complex dynamic BBB-NO co-culture model, the LOC device [17] was modified. Three organoid holders were integrated in the lower channel of the devices (Fig. 4A-B). The connected biochips received fresh media with an automatized syringe pump (Fig. 4A). The continuous fluid flow for dynamic conditions was maintained with a peristaltic pump in the upper channel. The LOC allowed the monitoring of the cell layer across the entire width and length of the membrane by microscopy. The human brain-like ECs showed characteristic F-actin and β-catenin staining patterns and the cell layer was confluent and intact (Fig. 4C).

**Fig. 4.**
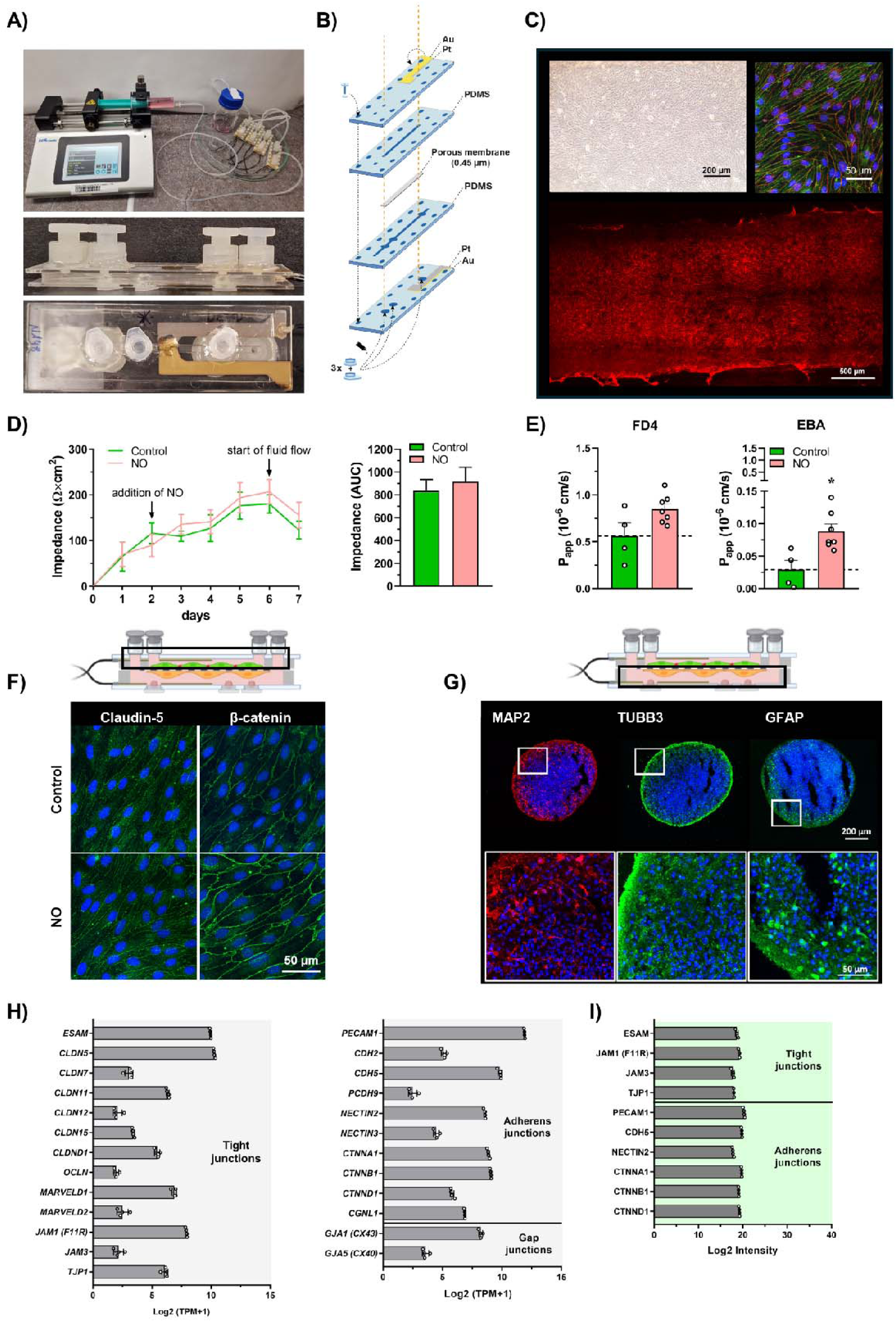
Characterization of the dynamic BBB-NO-LOC model. (A) Medium change with syringe pump on 4 connected lab-on-a-chip (LOC) devices; LOC from side and bottom view. (B) Schematic drawing of the LOC. The top and bottom layers are plastic slides with platinum-gold electrodes spin coated on their inner surfaces. The upper and lower PDMS channels are separated by a porous membrane. (C) Representative images of confluent human brain-like endothelial cell monolayers on the membrane. Top left panel: phase contrast microscopic image, scale bar = 50 µm. Top right panel: green: F-actin, red: β-catenin, blue: cell nucleus staining, scale bar = 50 µm. Bottom panel: red: β-catenin immunostaining on the whole endothelial cell layer, scale bar = 50 µm. (D) Kinetics of human brain-like endothelial cell growth in the LOC device with or without neural organoids (NO) measured by impedance for 7 days. AUC: area under the curve. Mean± SD, n=5-7, two-way ANOVA, Tukey and Bonferroni tests. (E) Dextran (FD4) and albumin (EBA) permeability on Day7. Unpaired t-test, Mean± SD, * *P*=0.0105, n=5-7. (F) Immunostaining of human brain-like endothelial cells. Green: claudin-5 and β-catenin, blue: cell nucleus staining, scale bar = 50 µm. (G) Immunostaining of neurons and astrocytes in neural organoids. Red: MAP2, Green: β3-tubulin and GFAP, blue: cell nucleus staining, scale bars = 200 µm and 50 µm. (H) Gene expression of cell-cell junction proteins (MACE-seq), n = 4. (I) Cell-cell junction protein abundance measured by proteomics. Mean± SD, n = 3.

#### Characterization of the BBB-NO-LOC model

Using the modified LOC device, we established a new dynamic BBB-NO co-culture model, which we characterized by morphological, functional, gene and protein expression analyses (Fig. 4-6). To establish the BBB model, human brain-like ECs and brain pericytes were seeded on the porous membrane (day 0), and 2 days later the neural organoids were added to the lower channel and were cultured together for 5 days. The impedance of the BBB model increased as the layer became confluent and the intercellular junctions tightened. The presence of brain organoids did not affect the tightness of the BBB (Fig. 4D). On day 6 we applied continuous fluid flow (1ml/min) for 24-hour in the upper channel that did not result in a statistically significant change in impedance values that exceeded 100 Ω×cm^2^ in both groups on day 7 (Fig. 4D). The apparent permeability coefficient for dextran was less than 1×10▯▯cm/s, for albumin less than 0.1×10 cm/s in both groups on day 7 (Fig. 4E), indicating good barrier function [19,27]. The morphology of human brain-like ECs in the BBB-NO-LOC model was characterized by immunocytochemistry for junctional proteins. Claudin-5 and β-catenin showed continuous belt-like pattern in both groups (Fig. 4F) and reflected the morphology of stem cell-derived human brain-like ECs from our previous studies [19,27]. These results show that the barrier properties of the BBB model remained unchanged in the presence of neural organoids.

The cellular morphology of the neural organoids after co-culture was characterized by immunostaining for neuronal markers MAP2 and β3-tubulin and astroglia marker GFAP. MAP2 labeled the processes and cell bodies of neurons and was uniformly distributed across the organoid surface, while β3-tubulin immunostaining outlined the periphery of the organoids. GFAP labeling was observed in clusters along the edges of the organoid (Fig. 4G). The morphology of neural organoids based on MAP2 staining was unchanged in the dynamic co-culture model (Fig. 4G) as compared to organoids kept without co-culture (Fig. 3D).

These data demonstrate that the BBB model can be co-cultured with neural organoids for several days and the barrier functions remain adequately tight (Fig. 4D-E). This co-culture setup was used for further experiments on the BBB-NO-LOC model.

The gene and protein expression profile in human brain-like ECs of the BBB-NO-LOC model was measured MACE-seq and proteomics (Fig. 4-6, Fig. S2-S4, Table S2). From the tight junction genes, *CLDN5*, *ESAM*, and *JAM1*, from the adherens junction genes *PECAM1*, *CDH5* coding VE-cadherin and *NECTIN2* showed the highest expression (Fig. 4H). The expression of ESAM, JAM1, PECAM1, VE-cadherin and NECTIN2 were verified by proteomics (Fig. 4I) and claudin-5 gave a continuous, belt-like junctional immunostaining (Fig. 4F). Among the cytoplasmic linkers the transcript and protein levels of ZO1 (*TJP1*) and catenins (*CTNNA1, -B1, -D1*) were also abundant. Gap junction genes *GJA1* and *GJA5* coding connexins 43 and 40 were also expressed (Fig. 4H). The presence of β-catenin was verified at protein level by both proteomics (Fig. 4I) and immunostaining (Fig. 4F). These key junctional genes and proteins were also present in the static contact [27] and dynamic non-contact [19] human BBB co-culture models characterized in detail in our previous publications.

The gene and protein expression of key endothelial markers VWF, CD34, NOS3, VEGFR2 in human brain-like ECs was high, validating the vascular endothelial nature of the model (Fig. 5A and 5G). Among the cell adhesion molecules ICAM-1 was present both at gene and protein levels, while the level of the inflammation-inducible *VCAM1* gene was low (Fig. 5B) and was not detected by proteomics (Fig. 5C). From the integrins ITGA1, ITGAV and ITGB5 (Fig. 5B and 5C), from the basal membrane proteins collagen type IV (COL4A2), fibronectin (FN1), perlecan (HSPG2), nidogen (NID1) and laminins (LAMA4 and 5) were detected by MACE-Seq (Fig. 5E) and proteomics (Fig. 5D). Important vesicular transport proteins caveolins (CAV1 and 2), cavins (CAVIN1-3), clathrins (CLTA, -B, -C), phosphatidylinositol binding clathrin assembly protein (PICALM) and osteonectin (SPARC) were well expressed (Fig. 5F and 5I) suggesting active endo- and transcytotic pathways. The gene and protein expression of additional vesicular transport associated proteins including adaptins, RABs, syntaxins, sorting nexins, cytoskeleton elements are listed in Table S2. The negatively charged glycocalyx is an essential physical barrier element at the BBB. Among the genes coding for endothelial cell surface glycocalyx proteins, *BGN*, *EXTL3*, *GPC1*, *PODXL*, syndecans *SDC3* and *SDC4*, *SPOCK2* and *TMEM123* were detected by MACE-Seq (Fig. 5J). The expression of podocalyxin was verified by proteomics (Fig. S2B). Several enzymes participating in glycocalyx synthesis and remodeling were detected at the gene level (Fig. S2A), from these HYAL2, HPSE and SULF2 were also detected at protein level (Fig. S2B).

**Fig. 5.**
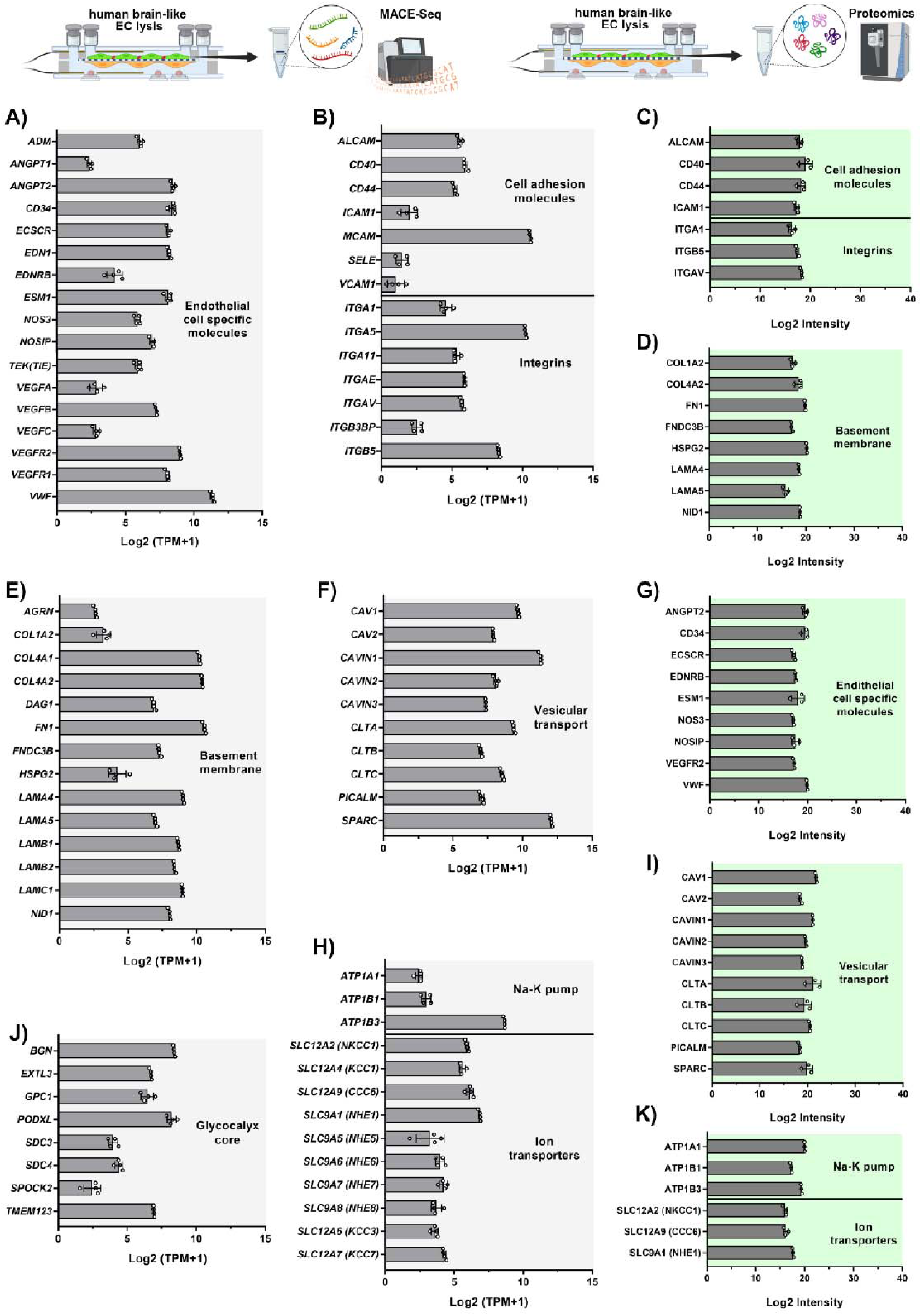
Expression of endothelial cell specific, cell surface, basement membrane proteins and ion transporters at gene- and protein level in cultured brain ECs in the dynamic BBB-NO-LOC model. (A) Endothelial cell specific genes. (B) Cell adhesion molecule and integrin genes. (C) Cell adhesion molecule and integrin proteins. (D) Basement membrane proteins. (E) Basement membrane genes. (F) Vesicular transport genes. (G) Endothelial specific proteins. (H) Na-K pump and ion transporter genes. (I) Vesicular transport proteins. (J) Glycocalyx core genes. (K) Na-K pump and transporter proteins. MACE-seq, n=4, TPM: transcript number/1 million transcripts; proteomics, n=3; mean ± SD.

The ATP1A1, 1B1, 1B3 subunits of the sodium pump, a BBB hallmark and a key transporter to provide ionic homeostasis in the CNS, is well expressed in the human brain-like human brain-like ECs, similarly to key ion transporters Na-K-Cl cotransporter NKCC1 (SLC12A2), cation-chloride cotransporter CCC6 (SLC12A9), and sodium-proton exchanger NHE1 (SLC9A1) (Fig. 5H and 5K). Ion channels for calcium, potassium and sodium expressed at gene level are shown in Fig. S3F. From the BBB carriers, transcripts were detected (Fig. 6A) for the following nutrient groups: hexose (*SLC2A1, 2A3*), cationic amino acids (*SLC7A1, 7A2, 7A6*), neutral amino acids (*SLC1A4, 1A5, 3A2, 6A6, 7A5, 38A1, 38A2*), anionic amino acids (*SLC7A11*), monocarboxylates (*SLC16A1, 16A3, 16A6*), organic cations (*SLC22A4, 22A5, 44A1*), organic anions (*SLCO2A1, O4A1*), fatty acids (*SLC27A1, 27A3, 27A4, 27A5, SLC59A1/MFSD2A*), vitamins and other small molecule nutrients (*SLC5A6, 23A2, 49A2*). From these carriers the expression of the BBB-specific glucose transporter-1 (SLC2A1), the amino acid transporter SLC1A5, the monocarboxylate transporter SLC16A3, the choline transporter-like protein 1 (SLC44A1), and the fatty acid transporters SLC27A3 and 27A4 was confirmed by proteomics (Fig. 6D). The BBB-specific efflux pump excitatory amino acid transporter-3 (*SLC1A1*) and multidrug and toxin extrusion protein 1 (*SLC47A1*) were also expressed at gene level (Fig. 6B). Among the ABC transporter transcripts the lipid transporter *ABCA3*, the drug efflux transporter P-glycoprotein (*ABCB1*), the multidrug resistance-associated proteins 1-6 (*ABCC1-6*) and the breast cancer resistance protein (*ABCG2*) were detected (Fig. 6B). From these, MRP1/ABCC1 was identified by non-targeted LC-MS based proteomics from cell lysates. The BBB expresses enzymes participating in local drug metabolism and efflux in coordination with ABC transporters. The phase-I drug metabolic cytochrome P450 enzymes CYP1A1 and CYP20A1, the phase-II epoxide hydrolase EPHX1, the BBB marker protein gamma-glutamyltransferase GGT5, glutathione S-transferases GSTK1, GSTKO1, MGST1 and MGST3 were detected by both transcriptomics (Fig. S2C) and proteomics (Fig. S2D). Among the BBB enriched receptors, the transferrin receptor 1 (TFRC), the immunoglobulin Fc receptor (FCGRT), the insulin-like growth factor 2 receptor (IGF2R), the scavenger receptor B2 (SCARB2) and the transmembrane protein 30A (TMEM30A), a phosphatidylserine flippase subunit, were expressed at both gene (Fig. 6C) and protein levels (Fig. 6E), while receptors for B12 vitamin (*CD320*), insulin (*INSR*), low density lipoprotein (*LDLR*), leptin (*LEPR*), apolipoprotein E (*LRP1*, *LRP8*), amyloid precursor protein (*LRP10*) and high density lipoprotein (*SCARB1*) were present at transcript level (Fig. 6C).

**Fig. 6.**
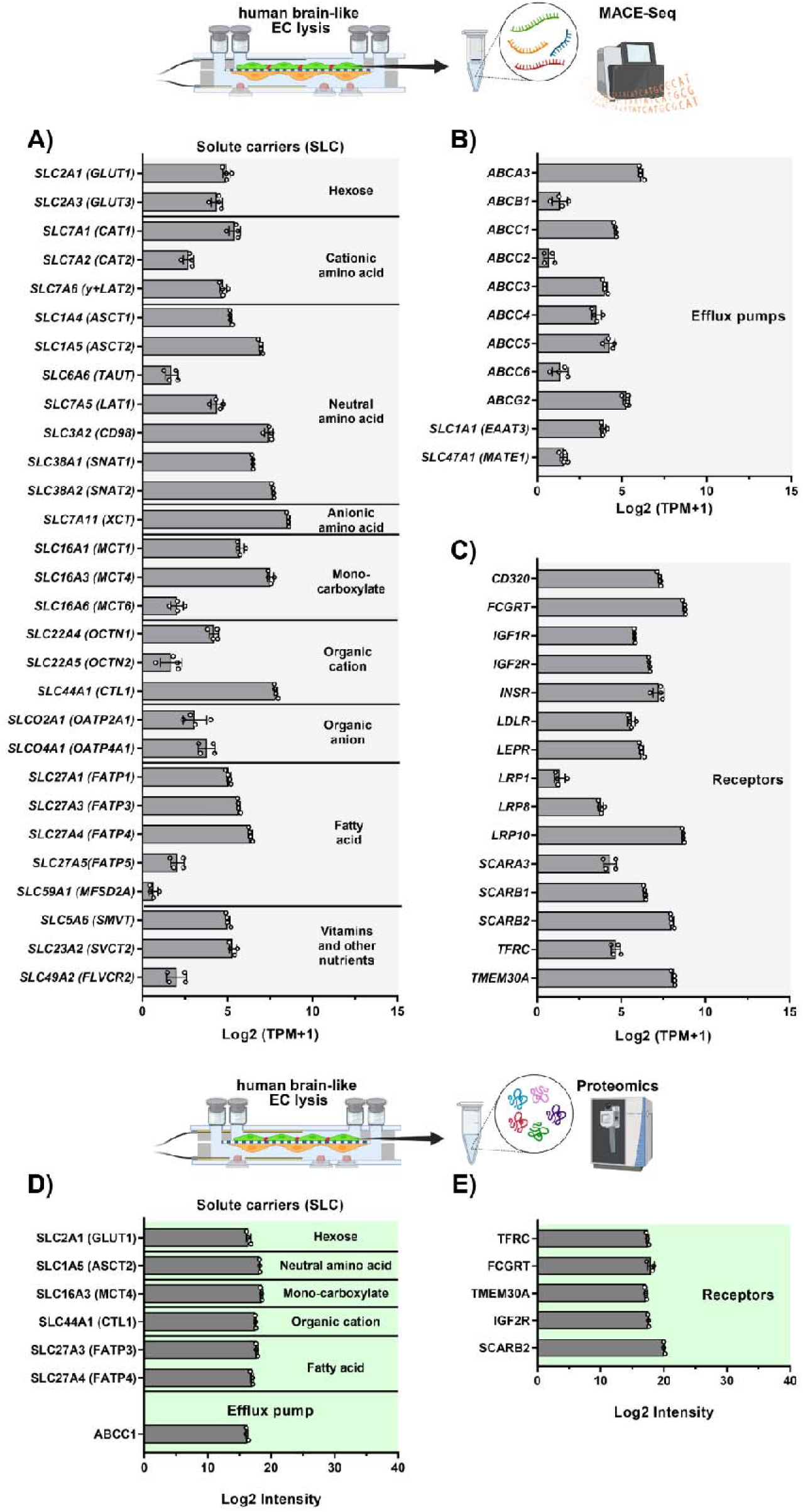
Expression of solute carriers, efflux pumps and receptors at gene- and protein level in cultured brain ECs in the dynamic BBB-NO-LOC model. (A) Solute carrier (SLC) genes. (B) Efflux pump genes. (C) Receptor genes. (D) SLC proteins. (E) Receptor proteins. MACE-seq, n=4, TPM: transcript number/1 million transcripts; proteomics, n=3; mean ± SD.

Endogenous glycan-binding proteins galectins (GAL1, 3, 8, 9) were well expressed in brain endothelial cells (Fig. S2E-F) together with the gene of collectin-12 (*COLEC12*), a C-type lectin scavenger receptor. The gene and protein expression of housekeeping proteins (Fig. S3A-B), metalloproteinases (Fig. S3C-D), endothelial transcription factors (Fig. S3E), endothelial cell surface mechanosensors, key molecules to transduce shear stress in endothelial cells, like *NOTCH1* and *PIEZO1* (Fig. S3G) and adhesion G protein-coupled receptors (Fig. S3H) are provided in the Supplementary data. To confirm that the medium composition does not affect significantly human brain-like ECs in dynamic conditions, we compared the transcriptome profile of the BBB-NO-LOC model compared to the BBB-LOC model without organoid, both cultured in the same culture medium (1:1 ratio of NO and EC media) as well as to BBB-LOC model cultured in EC medium (Fig. S4A-C). The Volcano plots showed no significant downregulated or upregulated genes.

#### Nanocarrier transport across the BBB-NO-LOC model

Nanocarriers are promising new delivery systems for enhancing drug delivery that have previously been studied by our group using static BBB models [23,36,37,38]. We studied two different nanocarriers, a polypeptide (Fig. 7B) and a vesicular nanoparticle (Fig. 7J) in the BBB-NO-LOC model in dynamic condition (Fig. 7A).

**Fig. 7.**
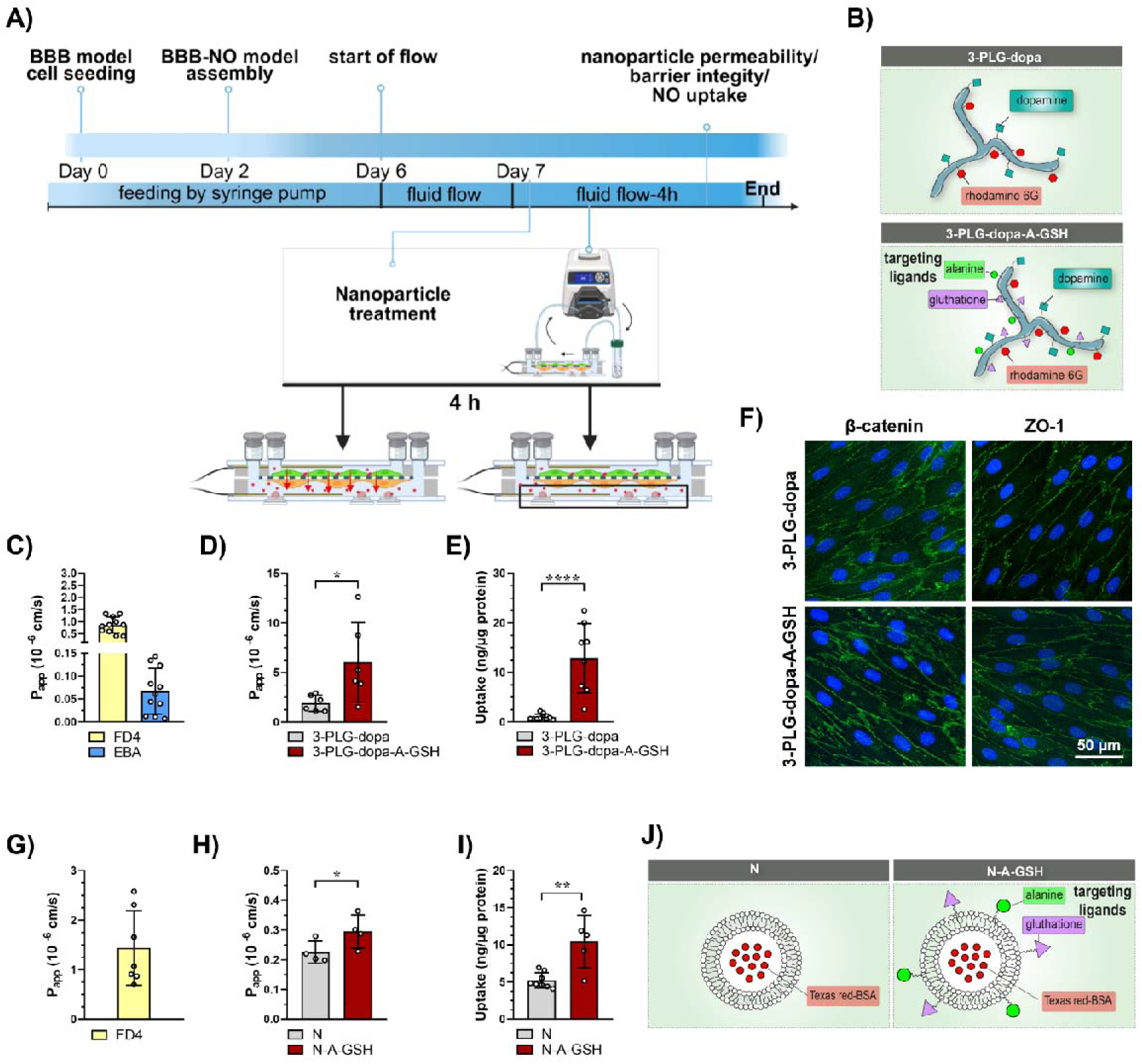
Polypeptide nanocarrier transport across the BBB and uptake into the neural organoids in the dynamic BBB-NO-LOC model. (A) Timeline of the experiment. (B) Schematic drawing of the structure of three-armed polypeptide nanocarriers with dopamine and rhodamine-6-G fluorescent labeling without targeting ligands (3-PLG-dopa), and with alanine and glutathione targeting ligands (3-PLG-dopa-A-GSH). (C) Apparent permeability coefficients (P_app_) for dextran (FD4) and albumin (EBA) across the BBB model. n = 11. (D) P_app_ of 3-PLG-dopa and 3-PLG-dopa-A-GSH nanocarriers, n = 6 chips/group. (E) Uptake of the nanocarriers into neural organoids (NO). Mean ± SD, unpaired t-test, * *P*<0.05, *** *P*<0.001, n = 8 organoids/group. (F) Immunostaining of tight junction proteins in brain endothelial cells. Green: β-catenin and ZO-1, blue: nuclei, scale bar = 50 µm. (G) P_app_ for FD4, n = 7 (H) P_app_ for alanin-glutation targeted vesicular nanocarrier (N-A-GSH) and non-targeted vesicular nanocarrier (N), n = 8. (I) Nanocarrier uptake in neural organoids. Mean ± SD, unpaired t-test, * p<0.05, ** p<0.01, n = 5-8. (J) Schematic drawing of the structure of targeted and non-targeted vesicular nanocarriers.

Dopamine was grafted on the three-armed poly(L-glutamic acid) nanocarrier as a therapeutic molecule which contained rhodamine 6G as a fluorescent label (3-PLG-dopa). Alanine and glutathione were used as targeting ligands (3-PLG-dopa-A-GSH; Fig. 7B). The baseline integrity of the BBB model was confirmed by dextran and albumin permeability, both of which remained low, consistent with a tight endothelial barrier (Fig. 7C). The targeted 3-PLG-dopa-A-GSH nanocarrier crossed the BBB model to a significantly greater extent than the non-targeted 3-PLG-dopa nanoparticle (Fig. 7D). A higher level of accumulation in neural organoids was also observed for the targeted nanoparticle (Fig. 7E). The immunostaining for β-catenin and ZO-1 in human brain-like ECs remained continuous in both groups indicating intact tight junction network after the assay (Fig. 7F).

The vesicular nanocarriers, the targeted and non-targeted niosomes (N and N-A-GSH) had comparable diameters (∼ 90 nm) and polydispersity index (∼ 0.25) (Table S3). We measured negative zeta potentials for both formulations (N: -8.53 ± 0.35 mV, N-A-GSH: -9.05 ± 0.07 mV) and for the Texas red-BSA cargo an encapsulation efficiency of 22.88% and 20.39%, respectively (Table S3). The biocompatibility of the fluorescently labelled cargo (Texas red-BSA) and the niosomes was evaluated by real-time impedance measurement on human brain-like ECs at various concentrations for 24 hours (Fig. S5). No effect on the human brain-like EC layer was observed for the Texas red-BSA cargo alone (1-10 µg/ml) during the experiment (Fig. S5A-B). Although a time- and concentration-dependent decrease in impedance was induced by the niosomes at the highest concentrations (6 and 10 mg/ml) and longer time points (Fig. S5C, S5E), at the 4-hour time point no significant change in cell index was measured for the non-targeted niosome (Fig. S5D), and only a 4% reduction in cell index was observed for N-A-GSH at the 3 mg/ml concentration (Fig. S5F). Based on these results, we selected the 3 mg/ml nanoparticle concentration and the 4-hour exposure time for further experiments.

As a further validation experiment the P_app_ of the niosomes, and the FD4 paracellular marker was compared on the static human BBB model cultured on membranes with 0.4 µm or 3 µm pore size. Both niosomes were found to cross the BBB model at a similar speed regardless of the membrane pore size, with no statistical difference detected between N and N-A-GSH in this setup (Fig. S5G). Low FD4 P_app_ (< 1.5 × 10 cm/s) was observed on the BBB model with both membrane types following the 4-hour nanoparticle assay, confirming that the barrier integrity remained intact (Fig. S5H).

In the dynamic BBB-NO-LOC model, the permeability of the N-A-GSH niosome was significantly higher than that of the N nanocarrier (Fig. 7H). Likewise, the entry of the N-A-GSH nanoparticles into the neural organoids was significantly increased compared to the uptake of the non-targeted ones (Fig. 7I). FD4 permeability across the dynamic BBB model following the 4-hour nanoparticle assay remained low (Fig. 7G), in a similar fashion to the static model (Fig. S5H), corroborating an unchanged barrier integrity.

The hyperosmolar iodinated contrast agent iopamidol (Fig. 8A) decreased the integrity of human brain-like ECs in a concentration- and time-dependent manner measured by cell impedance kinetics (Fig. S6A). Iopamidol exposure lasted for 30 minutes followed by cell culture medium change. Normalized cell index in the 10% iopamidol group was similar to the control group, while treatment with 50% and 100% iopamidol led to a significant drop in impedance, with the highest concentration (100%) showing no recovery over the 24-hour monitoring period. Quantification of the area under the curve confirmed the concentration-dependent decrease relative to the control group (Fig. S6A). For further experiments the 50% dilution of iopamidol was selected. On the human BBB model in static condition a 30-minute treatment with 50% iopamidol (acute treatment) significantly increased dextran permeability, which returned back to the level of the control group 24-hour after culture medium change (recovery group; Fig. S6B).

**Fig. 8.**
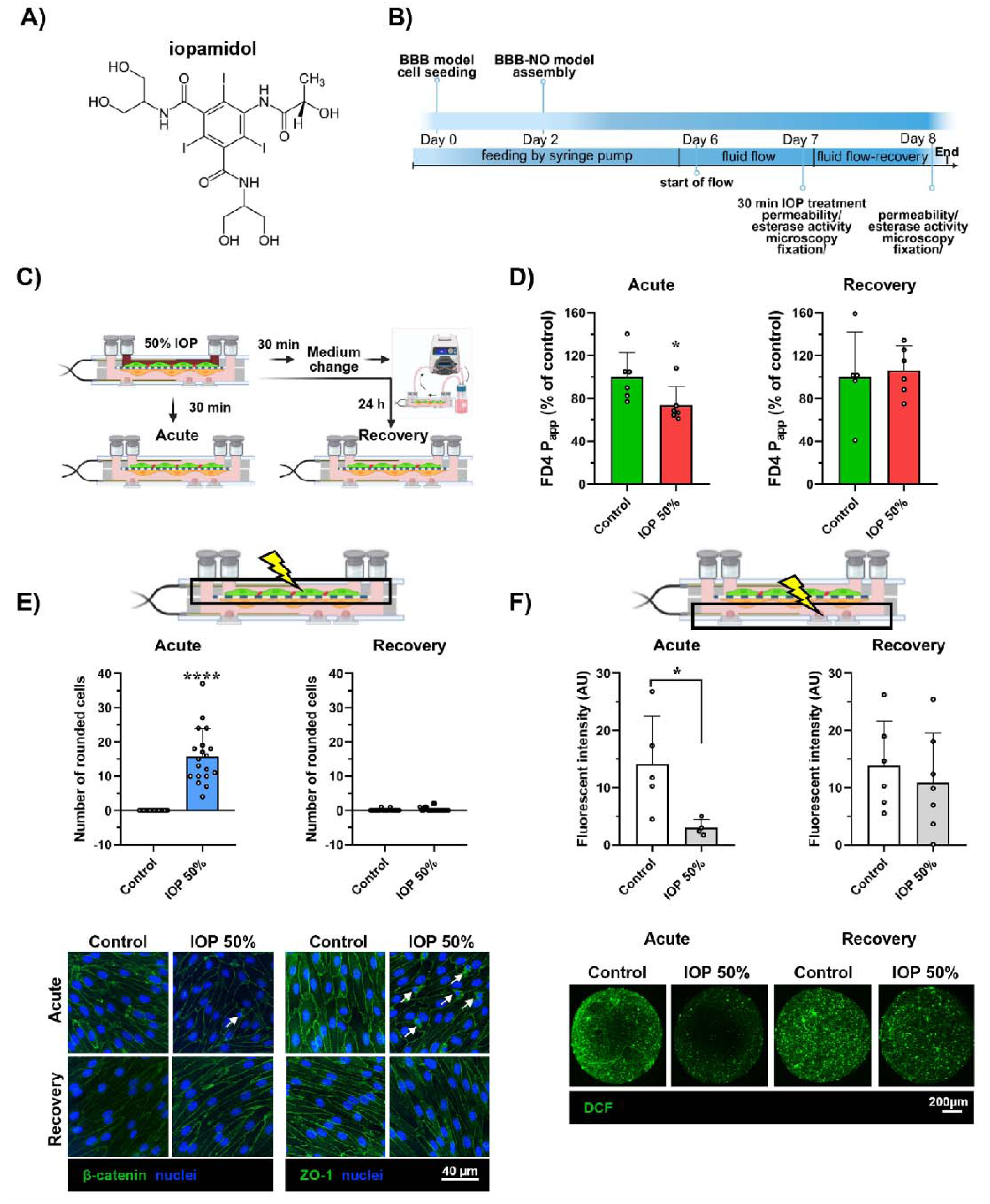
The effect of iopamidol contrast agent on the dynamic BBB-NO-LOC model. A) The structure of iopamidol (IOP) molecule. B) Experimental timeline. (C) Schematic drawing of the experimental groups. (D) Dextran (FD4) permeability assay after the 30-minute 50% IOP treatment (acute) or 24-hour after medium change (recovery). Mean ± SD, unpaired t-test, * *P*=0.0488, n = 5-6. (E) Number of human brain-like ECs detached after the 30-minute 50% IOP treatment (acute) or 24-hour after medium change (recovery). Mean ± SD, one-way ANOVA, **** P < 0.0001, n = 20. Green: β-catenin, blue: cell nuclei, scale bar = 40 µm. (F) Esterase activity measured by DCFDA dye conversion. Fluorescent intensity of neural organoids was measured right after the 30-minute 50% IOP treatment (acute) or 24-hour after medium change (recovery). Mean ± SD, unpaired t-test, * P = 0.0372, n = 4-7. Green: DCFDA, scale bar = 200 µm.

Direct iopamidol exposure also affected the metabolic function of neural organoids. The production of the fluorescent calcein from calcein AM by non-specific intracellular esterases decreased in a concentration-dependent way after 30-minute iopamidol treatment indicating a reduced metabolic activity (Fig. S6), which was confirmed by the diminished fluorescent signal intensity in the representative confocal microscopy images (Fig. S6D). After a 24-hour recovery period the metabolic activity of the organoids treated with the contrast agent was similar to the control group (Fig. S6C-D). Similar results were obtained when the fluorescence intensity values of neural organoids were normalized to protein content (Fig. S6E).

To evaluate the effects of the contrast agent on the barrier integrity of the dynamic BBB-NO-LOC model, the luminal surface of the brain-like endothelial cells was exposed to 50% iopamidol for 30 min (acute treatment), followed by a 24-hour recovery period in culture medium (Fig. 8B-C). Although the acute 50% iopamidol treatment did not increase the dextran P_app_ values as compared to control group (Fig. 8D), it significantly elevated the number of rounded endothelial cells in the process of detachment from the monolayer (Fig. 8E, arrows). This cell rounding as a step in detachment was not accompanied by visible disruption in the staining pattern of junctional linker proteins β-catenin and ZO-1 of the endothelial cells still forming monolayers. At the end of the 24-hour recovery period, the number of rounded endothelial cells returned to the level of the control group, with junctional staining patterns comparable between the control and iopamidol-treated groups (Fig. 8E). Neural organoids integrated in the dynamic BBB-NO-LOC model, were investigated by the DCFDA assay either in the acute iopamidol treatment phase or in the recovery phase (Fig. 8F). The fluorescent signal of the deacetylated and then oxidized intracellular DCF was significantly decreased after acute iopamidol treatment in neural organoids measured by fluorescent spectroscopy and this loss of signal intensity was well visible in the representative images, too. No change was seen between the groups after the 24-hour recovery period (Fig. 8F).

## Discussion

The first line of defense of the CNS is created by the brain barriers. The unique interaction between neurovascular cell types creates and maintains the BBB, a dynamic interface, where the brain endothelial cells form the anatomical basis [3]. With their specialized transporters these cells provide nutrients and oxygen to the brain, while restrict the entry of harmful substances. BBB dysfunction is related to many neurological and psychiatric diseases, but also to systemic and infectious conditions [1]. Brain endothelial cells and their tightly controlled transport functions, including efflux transporters, are mainly responsible for the low entry of a large group of pharmaceuticals to the CNS [6]. For these reasons, the creation and validation of complex, human cell-based cell culture models in biomedical research have never been more urgent.

The field of microphysiological systems and LOC modelling is getting more and more attention, significantly emerging in the past 15 years, and growing exponentially in the past 5 years [5]. The versatility of microfluidic and microelectronic devices along with the availability of human organoids, including neural organoids, provide unique tools for disease modelling, drug targeting and developmental studies [12,15,41,42]. To create LOC based microphysiological systems multidisciplinary collaboration is needed, which many research hubs lack [5]. The creation and differentiation of organoids to model different brain regions require such special expertise. With a consensus by leading researchers of the CNS organoid field, clear terms in nomenclature are set and being used for 3D model types, that we adapted in this paper. Neural organoids are created when various brain cells are differentiated from the same stem cell-origin and co-develop and self-organize with constant interaction [43].

In the present BBB-NO system a standardized and well characterized midbrain organoid model differentiated from iPSCs of a healthy donor was used. These spherical neural organoids contain neuronal and glial cells and do not develop a necrotic core [22]. The midbrain organoid model used in the present study was characterized in depth with single cell RNA sequencing and morphology studies [22,44,45]. When the human BBB co-culture model was integrated with neural organoids, BBB properties were stable both in the static system (Fig. 3; Fig. S1) and the dynamic model (Fig. 4). Both brain endothelial barrier integrity and intercellular junctional morphology were comparable to the control group. Since the NO and the EC culture media contain multiple growth factors or serum, it was crucial to establish that their optimal mixture at 1:1 ratio did not change tyrosine hydroxylase expression in neural organoids, and had no effect on the BBB properties, compared to the relevant control groups. The gene expression study showed that there was no difference in brain-like ECs when cultured in NO medium mixed with EC medium compared to EC medium alone (Fig. S4).

Static cell culture insert-based BBB models are the gold standard in *in vitro* brain microvascular research [6,46]. The development of co-culture models with multiple cell types [47], later the introduction of iPSC-derived [48,49,50] and hematopoietic stem-cell derived models [21] greatly advanced the field [6]. Our laboratory has been using the model established by Cecchelli et al. [21] in the last decade for multiple studies ranging from static and dynamic modelling, drug targeting by nanoparticles to investigations on BBB dysfunction. The stem-cell derived human vascular ECs when cultured together with brain pericytes develop brain-like properties with functional transporters and barrier integrity [51].

Dynamic flow in a previous version of our LOC device strengthened the BBB properties of brain-like ECs and increased the expression of glycocalyx components [19]. The modification of the LOC also enabled the measurement of the streaming potential of a brain endothelial monolayer [18]. In the present study our aim was to establish an even more complex system with the integration of the BBB co-culture model with neural organoids for biomedical investigations. The goal was to observe changes not only at the BBB level, but also at the neural organoid level representing the brain tissue compartment. In the BBB-NO-LOC system the BBB model is co-cultured with brain organoids forming a common niche during experiments, while enabling the independent analysis of all cellular and fluid compartments. The following aspects of the BBB-NO-LOC model make the system unique in its field: microfluidic LOC with active fluid flow created by a peristaltic pump, impedance measurement with Pt/Au electrodes, BBB co-culture model based on human stem cell derived ECs and brain pericytes integrated with iPSC-derived midbrain organoids (Fig. 9).

**Figure 9.**
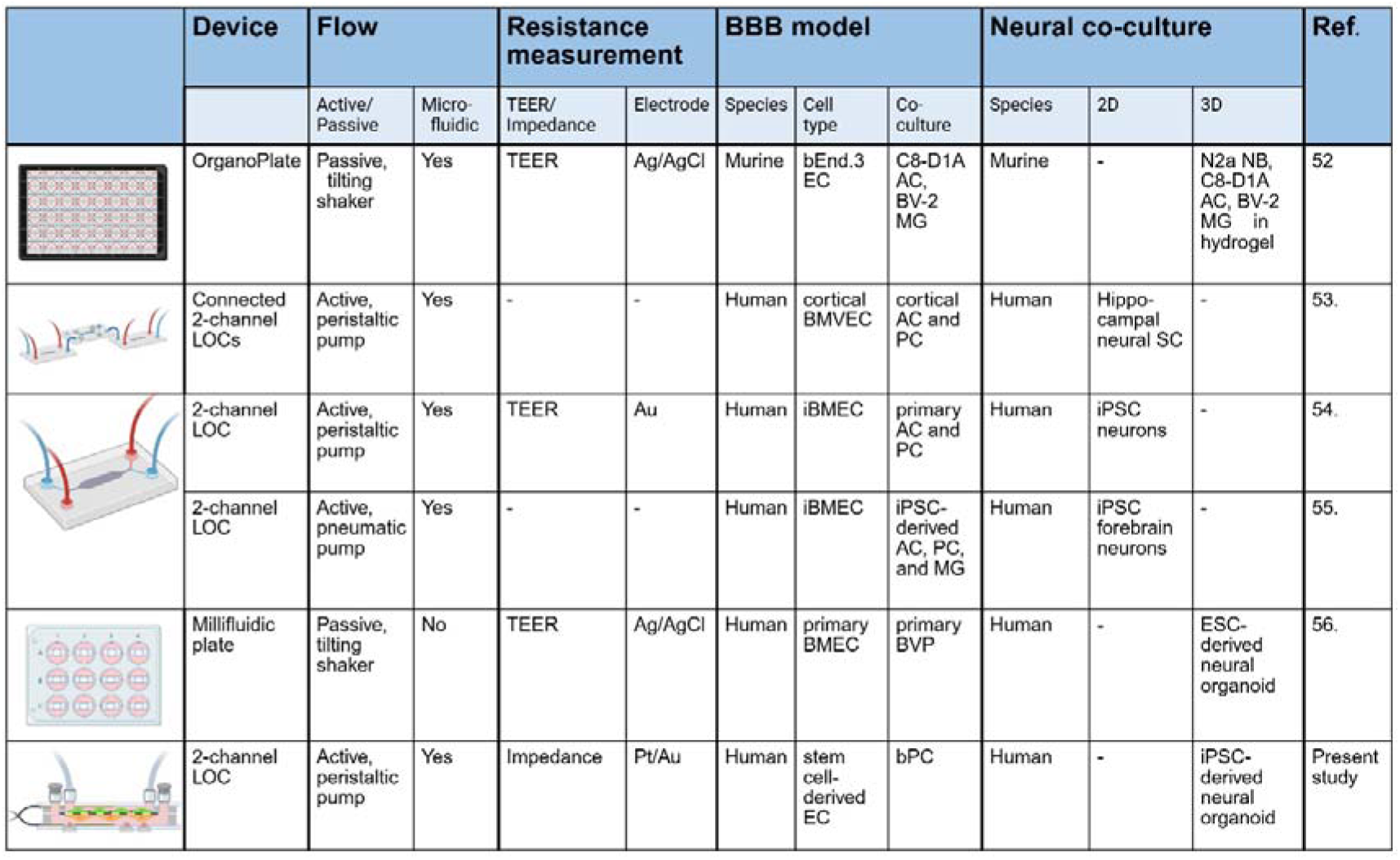
Summary of selected dynamic BBB models integrated with neural cultures. Abbreviations: AC: astrocyte; bEnd.3: immortalized murine brain endothelial cells; BMEC: brain microvascular endothelial cell; BMVEC: brain microvascular endothelial-like cells; bPC: bovine brain pericyte; BVP: brain vascular pericyte; EC: endothelial cell; ESC: embryonic stem cell; iBMEC: (iPSC)-derived brain microvascular endothelial-like cells; iPSC: induced pluripotent stem cell; LOC: lab-on-a-chip; MG: microglia; NB: neuroblastoma; PC: pericyte, SC: stem cell; TEER: transendothelial electrical resistance. Created in BioRender. Vigh, J. (2026) https://BioRender.com/zo00mt2

Creating vasculature in a neural organoid remains a challenge to the field. To date no vascularized brain cell type-based organoid model exists, where vessels could be perfused, or vasculature presents with mature and characterized BBB properties [14,57]. As a result, models integrating BBB and neural organoids in LOC may fill a gap as special tools in biomedical research. We selected those brain endothelial cell-based dynamic BBB systems that allow fluid flow, the measurement of barrier integrity by resistance or impedance, and the co-culture of multiple cell types, including neural cells. Studies using cancer cell types (glioma), hydrogel based self-organized brain microvascular models, or peripheral endothelial cells, such as HUVEC were not included. Multiple dynamic BBB systems were published previously, but in these 2D cultures of neurons [54,55], neurons and astrocytes [53], or a mixture of astrocytes, microglia cells and neurons in hydrogel (3D) were used for co-culture (Fig. 9). Except the BBB-NO-LOC model in the current work, only one study integrated a BBB co-culture model with neural organoids, but the device was a millifluidic plate with passive diffusion and not an LOC [56]. Importantly, targeted delivery across the BBB and entry into neural organoids was not investigated in these models.

Several advancements were made in our previously published LOC device [17,18,19] to integrate neural organoids. A novel Pt/Au electrode and removable holders were for the neural organoids were designed. These changes enabled the insertion and removal of three organoids in the bottom compartment of the device without disturbing the stability and integrity of the BBB co-culture. The previous unique features of the LOC, phase contrast and fluorescent microscopy on the whole culture surface, dynamic fluid flow, real time impedance and permeability measurements remained functional. All steps of the new design were carefully validated along with the static insert experiments. Impedance was followed for 7 days throughout the co-culture (Fig. 4). On day 7, the permeability values of paracellular markers indicated good barrier properties. Intercellular junction morphology in brain-like endothelial cells and neural organoid morphology based on neuronal and glial markers showed no alteration compared to the respective control groups reflecting a functional BBB and neural organoid niche.

To further characterize this unique model, the expression of key vascular endothelial and BBB specific genes was confirmed by MACE sequencing in human brain-like ECs cultured in the complex BBB-NO-LOC device, and we could confirm the presence of many of these molecules by proteomics or by immunostaining. Several cell-junction proteins previously described in brain endothelial cells including claudin-5, ESAM, JAM1 and JAM3, PECAM1, VE-cadherin, nectin-2, ZO-1, and catenins were detected at gene and protein levels. The vascular endothelial phenotype of the model was also supported by the expression of von Willebrand factor, VEGF receptor 2, CD34, endothelial nitric oxide synthase (NOS3), and angiopoietin-2. A brain endothelial cell-associated profile of integrins (ITGA1, ITGAV, ITGB5), basement-membrane proteins (type IV collagen, fibronectin, perlecan, and laminins), and adhesion molecules (ALCAM and ICAM-1) was also detected. The expression of caveolins, cavins, clathrin chains, osteonectin, and a long list of vesicular transport associated molecules indicate that endocytic and transcytotic pathways are active in the model.

In addition to the classical BBB marker Na^+^/K^+^-ATPase, the expression of ion transporters (NKCC1, NHE1, and CCC6) was detected. Transport proteins for major nutrient classes were also identified, including glucose transporter 1; the neutral amino acid transporter ASCT2/SLC1A5; the monocarboxylate transporter MCT4/SLC16A3; choline transporter-like protein 1 (CTL1/SLC44A1); fatty-acid transporters FATP3/SLC27A3 and FATP4/SLC27A4; and the efflux transporter MRP1/ABCC1. The BBB-associated gene-expression pattern detected in human brain-like ECs of the dynamic BBB-NO-LOC model was similar to that reported previously for the same BBB model [19,27].

A greater number of BBB-specific genes were detected by sequencing than by proteomic analysis. This difference may be explained, in part, by the lower sensitivity of the untargeted proteomic method used in this study compared with MACE-seq. In addition, many cell-junction, adhesion, and transporter proteins are localized to the brain endothelial cell membranes, particularly within membrane microdomains with specialized lipid compositions [58]. Because a separate membrane fraction was not isolated before proteomic analysis, enrichment and subsequent detection of these proteins may have been limited.

For the investigation of targeted nanocarrier transport across a barrier model, the presence of the relevant target proteins, such as receptors, transporters, and surface molecules, need to be verified. In the present experiments, nanocarrier targeting was achieved using a combination of the amino acid alanine and the tripeptide glutathione based on our previous results [37,38,59]. In the brain-like ECs, the expression of neutral amino acid transporter genes involved in alanine transport, including SNAT2/SLC38A2, ASCT2/SLC1A5, ASCT1/SLC1A4, and SNAT1/SLC38A1, were detected. A similar expression profile had previously been reported in primary rat brain endothelial cells and isolated rat brain capillaries [37]. Glutathione transport across the plasma membrane has been associated with OATP and MRP transporters [60]. In the present model, gene expression of the SLC transporters OATP2A1 and OATP4A1 and of the ABC transporters MRP1-6 were confirmed. The presence of ASCT2 and MRP1 were also verified by proteomic analysis.

We have previously investigated and characterized the entry of A-GSH-targeted nanoparticles into brain organoids [23,59]. In these previous experiments the BBB model was kept together with neural organoids in a static cell culture insert setup and only for 24 hours during the experiment. Here the goal was to validate our dynamic BBB-NO-LOC system by a proof-of-concept experiment, showing that nanocarriers that pass through the brain endothelial cell layer are able to enter neural organoids representing brain tissue in the present model. A greater transport across the human brain-like EC layer and an increased entry into neural organoids were observed for both the targeted polymeric and vesicular nanocarriers compared to non-targeted ones. The present observations were similar to the results of previous studies from our laboratory in static BBB models [37,38,59], providing foundational data corroborating the usability of the present chip our system. Collectively, these data support the conclusion that dual targeting with alanine and GSH may enhance the transport of nanocarriers across the BBB and facilitate their entry into the brain and corroborate the applicability of the dynamic human BBB-NO-LOC model for nanoparticle experiments.

Another fundamental aim was to prove the usability of the novel BBB-NO-LOC system to answer translational biomedical questions. Contrast agents are widely used for CNS imaging and during neurovascular interventions. Despite neurological complications such as contrast-induced encephalopathy [61] little research has been performed to reveal the underlying mechanism and the direct effect of contrast media on the integrity of the BBB. In a recent study our teams have shown that acute treatment with iopamidol increased brain endothelial cell permeability in a rat primary triple co-culture BBB model [40]. The changes returned to level of the control group within 24 hours, in a similar way to the results of the current static human co-culture BBB experiments. Although 50% dilution of iopamidol did not increase the permeability of the endothelial layer in the dynamic BBB-NO-LOC system, morphological analysis showed an increase in rounded endothelial cells in the process of detachment from the EC monolayer, a well-characterized step in apoptosis [62]. The unique design of the dynamic BBB-NO-LOC system helped us to reveal that iopamidol given in the luminal compartment, representing the blood, induced in neural organoids a decrease in metabolic activity similar to when treated directly. This is in line with a metabolic-stress related chain reaction underlying iodinated contrast media induced neurotoxicity [63]. The phenomenon indicates that although the regeneration and recovery of the BBB may be rapid after cellular stress induced by hyperosmolar iodinated contrast agent, it can still cause neural organoid injury. The BBB-NO-LOC device does not only enable the separate observation of phenomena at the BBB or neural organoid level, but provides a platform where brain endothelial dysfunction can directly influence metabolic activity, viability, and other functional changes at the neural organoid level.

## Conclusion

Here we present a novel, dynamic, human cell-type based BBB-NO model in a LOC device, where continuous monitoring of barrier properties is possible. Barrier and other brain endothelial cell functions, morphology and molecular traits are stable during 5-day co-culture with neural organoids. Endothelial identity and BBB characteristics of brain endothelial cells in the BBB-NO-LOC were proven by gene sequencing and proteomics. The model is suitable to study targeted nanocarrier penetration across the BBB and entry into neural organoids, as well as to investigate the complex effect of clinically used molecules, such as hyperosmotic contrast agents, on both BBB integrity and neuronal metabolism. The BBB-NO-LOC device opens a new path for integrated lab-on-a-chip devices, where vessel and neural properties can be simultaneously investigated with two-way interaction between compartments. This leads to the possibility to study the effects of neural dysfunction on healthy BBB and vice versa. The present BBB-NO-LOC model was assembled by using stem cell-derived human brain endothelial cells and NOs from healthy donor iPSCs. In future experiments, the use of BBB models and neural organoids from disease specific iPSC in the LOC device could lead to a more complex investigation of diseases of the CNS and contribute to the comparison of potential therapeutics for personalized medicine.

## CRediT authorship contribution statement

Conceptualization: J.P.V., A.E.K., F.R.W., A.D., M.A.D.; Formal analysis: J.P.V., A.E.K., A.R.S.M., E.B., S.B., A.K., A.S., M.M., G.P., R.W., Z.S., M.A.D., F.R.W. Methodology: J.P.V., A.E.K., A.R.S.M., N.K., S.B., J.C.S., A.K., S.V., A.D., M.A.D., F.R.W., Software: E.B., A.K., R.W., Validation: J.P.V., A.E.K., A.R.S.M., F.R.W., Investigation: J.P.V., A.E.K., A.R.S.M., N.K., E.B., S.B., A.K., S.V., A.S., Sz.V., M.M., G.P., T.H.M.P., Z.S., F.R.W. Resources: J.C.S., Sz.V., T.H.M.P., J.S.J., M.C., Y.M., Data Curation: J.P.V., E.B., Writing – original draft: J.P.V., A.E.K., E.B., F.R.W., M.A.D.; Writing – Review and editing: J.P.V., A.E.K., A.R.S.M., N.K., E.B., S.B., J.C.S., S.V., A.S., Sz.V., M.M., G.P., T.H.M.P., J.S.J., R.W., Z.S., M.C., Y.M., A.D., M.A.D., F.R.W., Visualization: J.P.V., A.E.K., N.K., E.B., Supervision: F.R.W., M.A.D. Project administration: J.P.V., F.R.W., M.A.D. Funding acquisition: M.A.D., F.R.W. All authors reviewed, edited, and approved the final manuscript.

**Judit P. Vigh:** Conceptualization, Formal Analysis, Methodology, Validation, Investigation, Data Curation, Writing – original draft, Writing – Review and editing, Visualization, Project administration

**Anna E. Kocsis:** Conceptualization, Formal Analysis, Methodology, Validation, Investigation, Writing – original draft, Writing – Review and editing, Visualization

**Ana R. Santa-Maria:** Formal Analysis, Methodology, Validation, Investigation, Writing – Review and editing

**Nóra Kucsápszky:** Methodology, Investigation, Writing – Review and editing, Visualization

**Emese Bató:** Formal Analysis, Software, Data Curation, Writing – original draft, Writing – Review and editing, Visualization

**Silvia Bolognin:** Formal Analysis, Methodology, Investigation, Writing – Review and editing

**Jens C. Schwamborn:** Resources, Methodology, Writing – Review and editing, Funding acquisition

**András Kincses:** Formal Analysis, Software, Investigation, Methodology

**Sándor Valkai:** Methodology, investigation, Writing – Review and editing

**Anikó Szecskó:** Formal Analysis, Investigation, Writing – Review and editing

**Szilvia Veszelka:** Investigation, Writing – Review and editing

**Mária Mészáros:** Formal Analysis, Investigation, Writing – Review and editing

**Gerg**ő **Porkoláb:** Formal Analysis, Investigation, Writing – Review and editing

**Thi Ha My Phan:** Investigation, Resources, Writing – Review and editing

**Jeng-Shiung Jan:** Investigation, Resources, Writing – Review and editing

**Roland Wirth:** Formal Analysis, Software, Writing – Review and editing

**Zoltán Szabó:** Formal Analysis, Investigation, Writing – Review and editing

**Maxime Culot:** Resources, Writing – Review and editing

**Yoichi Morofuji:** Resources, Writing – Review and editing

**András Dér:** Conceptualization, Methodology, Writing – Review and editing

**Mária A. Deli:** Conceptualization, Formal Analysis, Methodology, Writing – original draft, Writing – Review and editing, Supervision, Project Administration, Funding acquisition

**Fruzsina R. Walter:** Conceptualization, Formal Analysis, Methodology, Validation, Investigation, Writing – original draft, Writing – Review and editing, Supervision, Project Administration, Funding acquisition

## Acknowledgments

Authors acknowledge the support of Gábor Steinbach and the Laboratory of Cellular Imaging, Complex Molecular and Cell Biology Service Centre, HUN-REN Biological Research Centre, Szeged, Hungary. We thank Tibor Páli (Institute of Biophysics, HUN-REN BRC) for measurements on a Horiba Jobin-Yvon Fluorolog 3 spectrofluorometer. We are grateful to GenXPro GmbH (Frankfurt, Germany) for the help with MACE-seq analysis and Ana Martins for the critical reading of the manuscript. We acknowledge the support of Péter Galajda (Institute of Biophysics, HUN-REN BRC) for providing the clean room facilities to all of our fabrication experiments.

## Appendix A. Supplementary data

Supplementary data to this article can be found online at the journal’s homepage.

## Data availability

The generated and analyzed MACE-seq data have been deposited to the Gene Expression Omnibus (GEO) repository under accession number (link will be provided after publication or private request).

All data needed to evaluate the conclusions in the paper are present in the paper and/or the Supplementary Materials are available upon request.

## SUPPLEMENTARY Materials

**Table S1.**
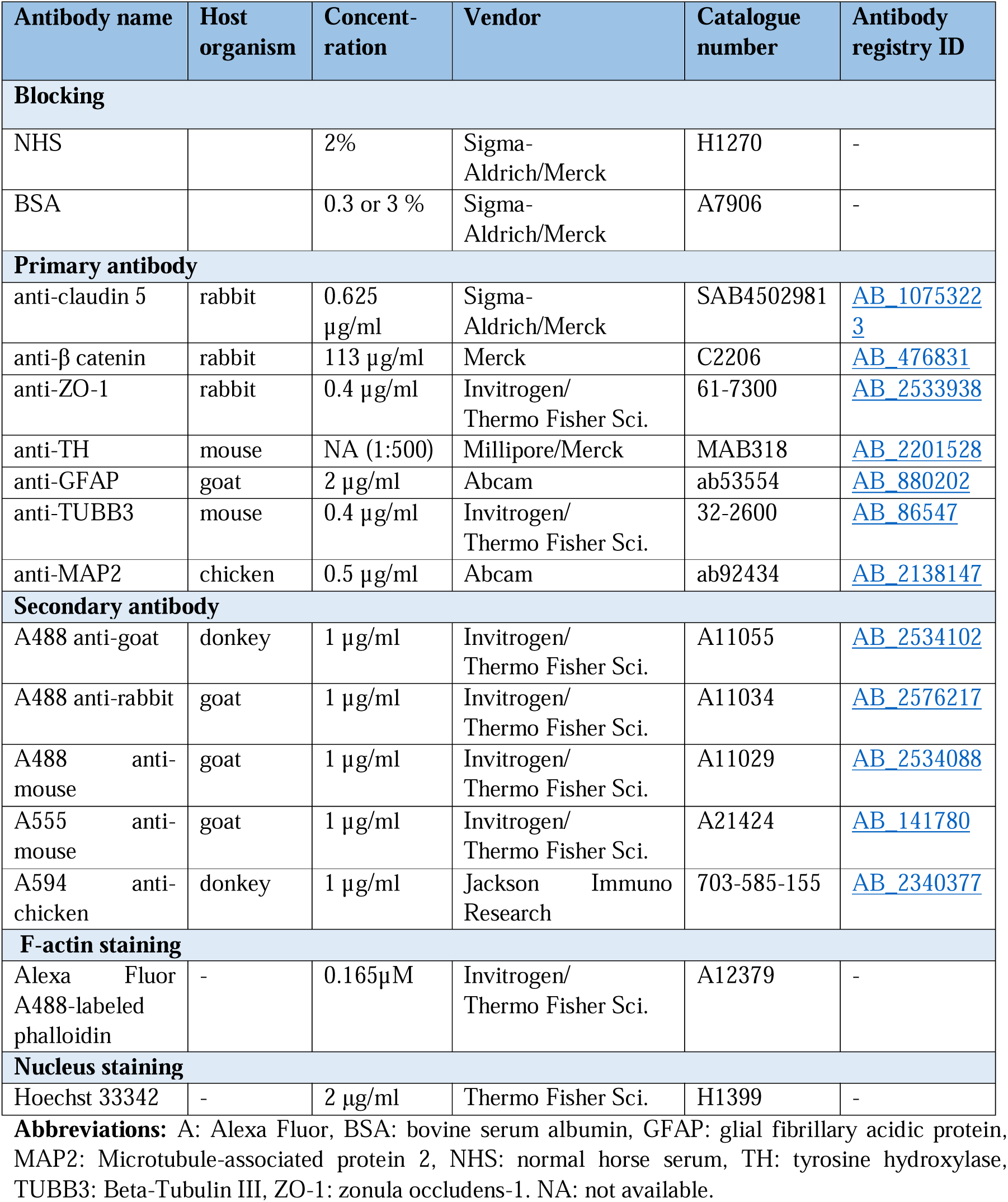
List of antibodies and solutions used for immunostaining.

| Antibody name | Host organism | Concentration | Vendor | Catalogue number | Antibody registry ID |
| --- | --- | --- | --- | --- | --- |
| **Blocking** |  |  |  |  |  |
| NHS |  | 2% | Sigma-Aldrich/Merck | H1270 | - |
| BSA |  | 0.3 or 3 % | Sigma-Aldrich/Merck | A7906 | - |
| **Primary antibody** |  |  |  |  |  |
| anti-claudin 5 | rabbit | 0.625 µg/ml | Sigma-Aldrich/Merck | SAB4502981 | AB_10753223 |
| anti-β catenin | rabbit | 113 µg/ml | Merck | C2206 | AB_476831 |
| anti-ZO-1 | rabbit | 0.4 µg/ml | Invitrogen/Thermo Fisher Sci. | 61-7300 | AB_2533938 |
| anti-TH | mouse | NA (1:500) | Millipore/Merck | MAB318 | AB_2201528 |
| anti-GFAP | goat | 2 µg/ml | Abcam | ab53554 | AB_880202 |
| anti-TUBB3 | mouse | 0.4 µg/ml | Invitrogen/Thermo Fisher Sci. | 32-2600 | AB_86547 |
| anti-MAP2 | chicken | 0.5 µg/ml | Abcam | ab92434 | AB_2138147 |
| **Secondary antibody** |  |  |  |  |  |
| A488 anti-goat | donkey | 1 µg/ml | Invitrogen/Thermo Fisher Sci. | A11055 | AB_2534102 |
| A488 anti-rabbit | goat | 1 µg/ml | Invitrogen/Thermo Fisher Sci. | A11034 | AB_2576217 |
| A488 anti-mouse | goat | 1 µg/ml | Invitrogen/Thermo Fisher Sci. | A11029 | AB_2534088 |
| A555 anti-mouse | goat | 1 µg/ml | Invitrogen/Thermo Fisher Sci. | A21424 | AB_141780 |
| A594 anti-chicken | donkey | 1 µg/ml | Jackson Immuno Research | 703-585-155 | AB_2340377 |
| **F-actin staining** |  |  |  |  |  |
| Alexa Fluor A488-labeled phalloidin | - | 0.165µM | Invitrogen/Thermo Fisher Sci. | A12379 | - |
| **Nucleus staining** |  |  |  |  |  |
| Hoechst 33342 | - | 2 µg/ml | Thermo Fisher Sci. | H1399 | - |

**Abbreviations:** A: Alexa Fluor, BSA: bovine serum albumin, GFAP: glial fibrillary acidic protein, MAP2: Microtubule-associated protein 2, NHS: normal horse serum, TH: tyrosine hydroxylase, TUBB3: Beta-Tubulin III, ZO-1: zonula occludens-1. NA: not available.

### Proof-of-concept functional tests

#### Nanoparticle penetration across the BBB and into the neural organoids

##### Preparation of nanovesicles

Vesicular nanoparticles (NPs) were made from non-ionic surfactants and cholesterol. For the preparation of non-tagged nanovesicles (N), Span 60 (sorbitane-monostearate), Solulan C24 (cholesterylpoly-24-oyxyethylene-ether, Chemron Co., USA) and cholesterol were dissolved in hot 1:2 mixtures of chloroform and ethanol in a round-bottom flask [1,2,3,4]. In the case of targeted nanoparticles, DSPE-PEG-linked alanine and glutathione (N-A-GSH; 4.5% w/w of total lipids) were added to the mixture of nanoparticle materials, and the solvent was removed under vacuum to yield a thin lipid film. The dry lipid film was hydrated with PBS containing 0.2 mg/ml Texas red-labeled bovine serum albumin (TR-BSA, 67.12 kDa, Thermo Fisher Scientific, USA) as cargo. The mixture was heated to 45 °C in a water bath and sonicated for 1 h. The suspension was filtered through a syringe filter with 0.45 μm pore size (Sarstedt, Germany) to yield vesicles. The non-entrapped cargo was removed by ultracentrifugation (123,249 *g*, 3 h, 4 °C), which resulted in a supernatant and a wet pellet on the bottom of the tube. The wet pellet contains the hydrated lipid bilayers that form the nanoparticles. After the aqueous supernatant phase was carefully removed by a micropipette, the mass of the remaining wet pellet was measured. To prepare a 100 mg/ml concentration of NPs the pellet was resuspended in the required amount of gentamycin-containing, phenol red-free DMEM/HAM’s F-12 culture medium (Gibco, USA). Nanoparticles were filtered with 0.2 μm pore size syringe filter (Sarstedt, Germany) and stored at 4 °C until further experiments.

#### Characterization of nanoparticles

##### Size, polydispersity index and surface charge measurements

To characterize the physico-chemical properties, NPs were diluted in PBS or distilled water to a final concentration of 5 mg/ml. The hydrodynamic size, polydispersity index (PDI) and surface charge (zeta potential) of NPs were measured by dynamic light scattering (DLS) (Malvern Zetasizer Ultra, UK).

##### Encapsulation efficiency of NPs (EE %)

The amount of encapsulated cargo in the synthesized NPs was determined by indirect method [3,4]. Briefly, the non-entrapped TR-BSA cargo was detected in the supernatant from the last ultracentrifugation step, by spectrofluorimetry (Fluorog 3, Horiba Jobin Yvon, France) at 589/609 nm. The encapsulation efficiency percentage (EE %) was calculated by the following equation:

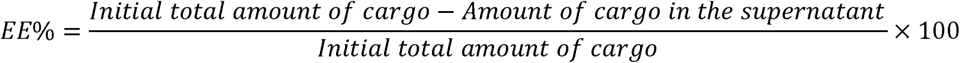

##### Measurement of cellular toxicity

Kinetics of the effect of NPs on human brain-like EC cells was monitored by real-time cell impedance measurement in a multiwell plate, treated after human brain-like ECs reached the plateau phase, similarly as described in the main text. Here cells were treated with N-A-GSH-tagged and non-tagged NPs (1-10 mg/ml) for 24 h. Impedance was followed for 24 h.

##### Permeability studies on static BBB models

For permeability studies, the contact co-culture BBB model was used, in which human brain-like EC and bPC are cultured together in a cell culture insert, similarly as described in the main text “Static cell culture model”, just without neural organoids. Here two types of cell culture inserts were used: polyethylene terephthalate (PET) membranes, 0.4 and 3 μm pore sizes (Corning Costar, USA) coated with collagen type IV (100 μg/ ml). The BBB model was cultured together for one week before permeability measurements in culture media.

To test the penetration of targeted and non-tagged NPs, 3 mg/ ml NPs were added to the the donor compartment of the BBB model for 4 h, diluted in phenol red-free DMEM/HAM’s F-12 medium supplemented with 5% FBS and 100x ECGS. After NP permeability, to assess the integrity of the model, the passage of the paracellular permeability marker, the 4 kDa FITC-dextran (FD4; 100µg/ml) was also tested for 30 min. After the permeability assays samples were collected from both compartments and the fluorescent signal of TR-BSA (excitation: 592 nm; emission: 613 nm) and FD4 marker (excitation: 490 nm; emission: 516 nm) were quantified with a spectrofluorometer (Horiba Jobin Yvon Fluorolog 3, USA). For the determination of permeability, the apparent permeability coefficients (P_app_) were calculated as described in the main text.

#### Testing the effects of a hyperosmolar contrast agent on human brain-like endothelial cells and neural organoids

The effects of iopamidol were pre-tested before the experiments on the LOC device by using static cell culture inserts. The model was assembled similarly to the static insert model using human brain-like EC and brain pericyte contact co-culture, just without neural organoids. Cells were treated for 30 min with 50% iopamidol, then permeability of FD4 was tested. Treatment solution was replaced in the recovery group with human brain-like EC cell culture medium, and FD4 penetration was tested after 24 h. Papp was calculated after the quantification of the FD4 fluorescent signal from the bottom compartment as described above and in the main text.

Neural organoids were cultured in low-attachment 96-well plates (Corning) and treated directly for 30 min. Metabolic activity of “acute” and “recovery” groups was tested by Calcein AM assay, similarly as described in the main text. For the Calcein AM, representative pictures of neural organoids was also taken with a Leica TCS SP5 confocal laser scanning microscope to show the enzymatic activity and its recovery in each group.

**Fig. S1.**
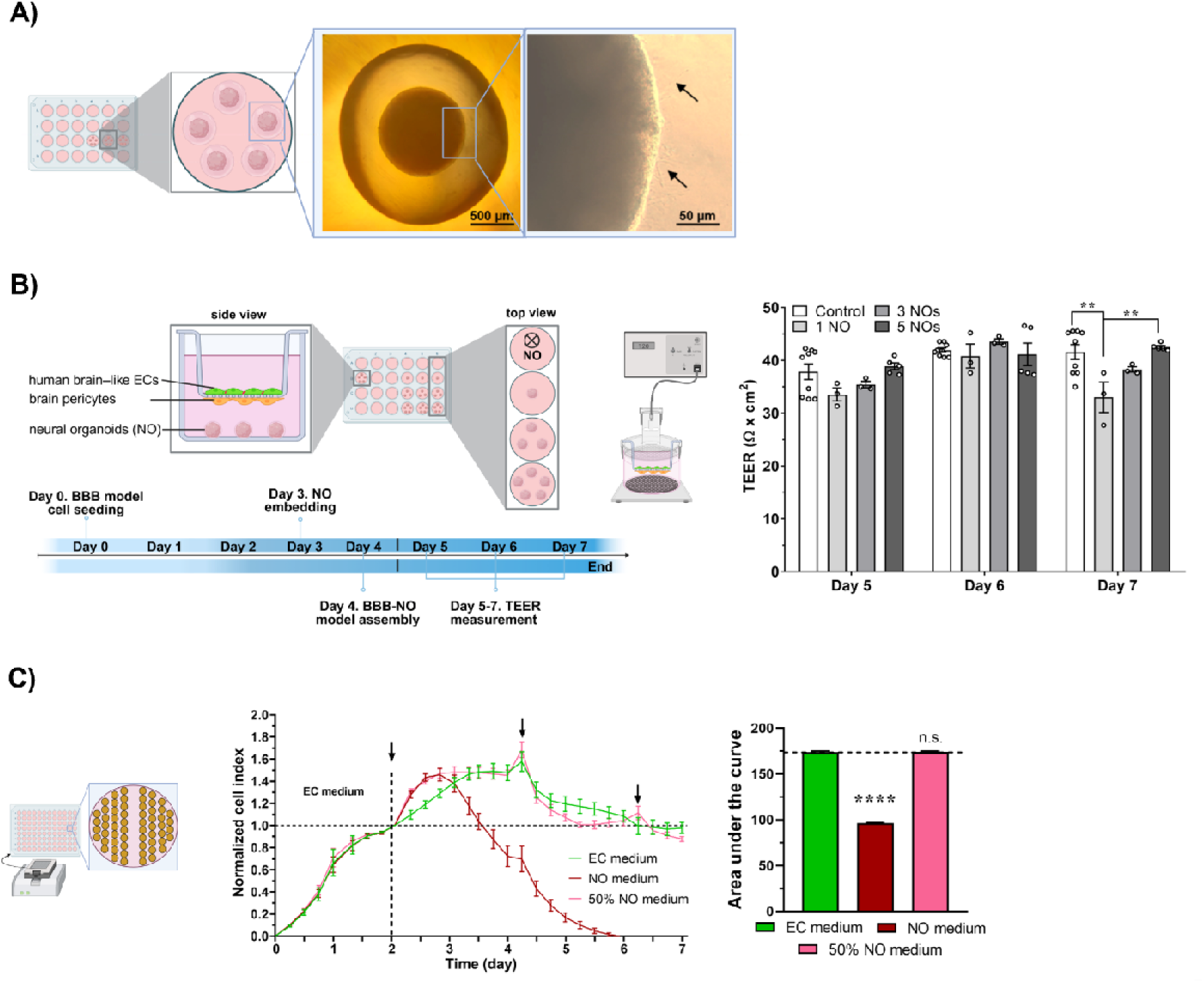
Optimization of the static BBB-NO co-culture model. (A) Schematic drawing and phase contrast image of neural organoids after their embedding in Matrigel. Arrow: neurite outgrows. Scale bars: 500 µm and 50 µm. (B) Effect of the number of neural organoids on the contact brain EC-pericyte co-culture BBB model. Schematic drawing of the experimental setup and timeline or the assembly and co-culture. Transendothelial electrical resistance (TEER) measurement was performed between Day 5-7. Two-way ANOVA, Bonferroni post-test, mean ± SD, n = 3-9, ** P<0.01. (C) Real time cell impedance analysis of human brain-like EC monolayer. NO medium treatment with different concentrations (50% and 100%). One-way ANOVA, Bonferroni post-test, mean ± SD, n = 8, **** *P*<0.0001.

**Fig. S2.**
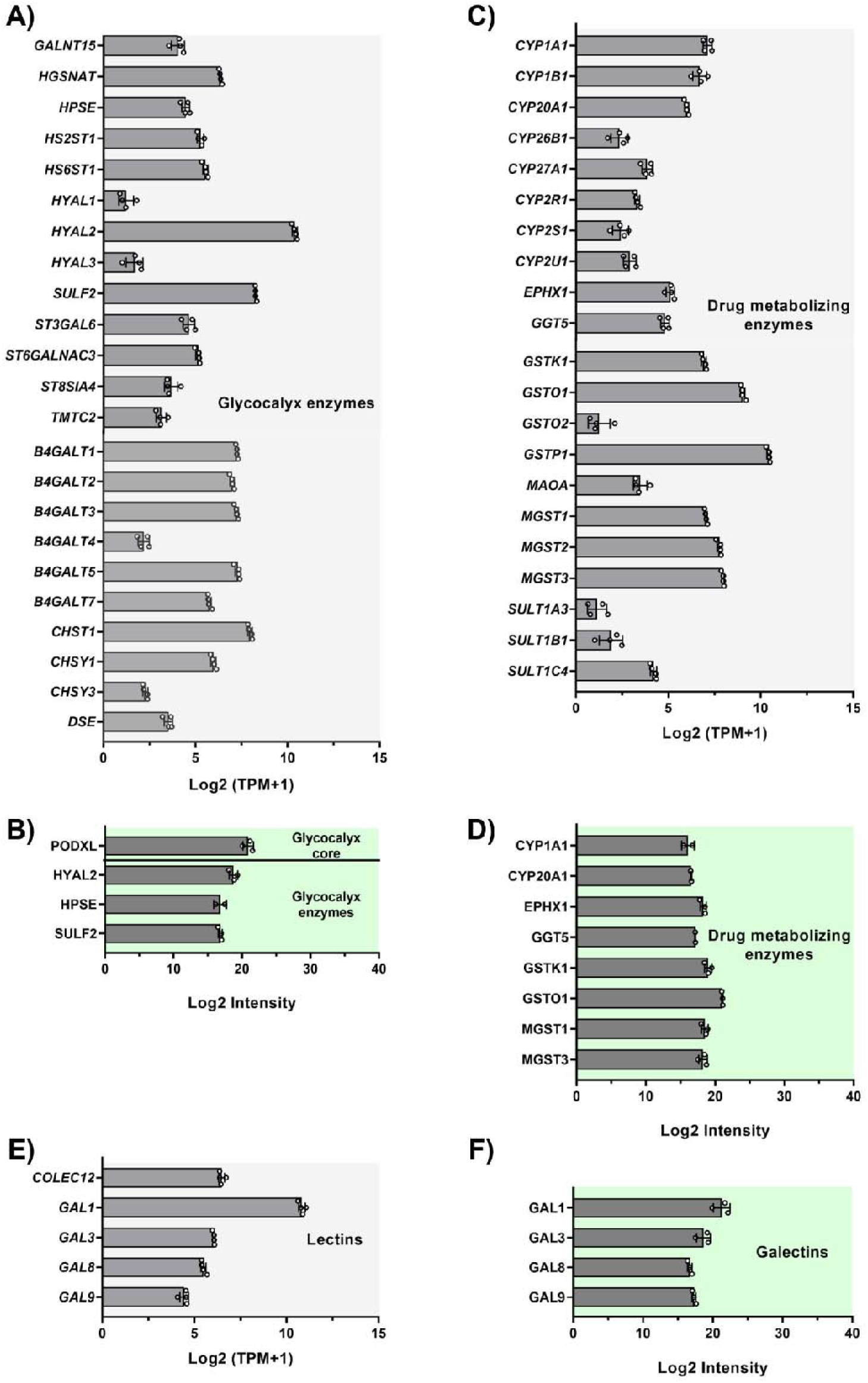
Gene- and protein expression of cultured brain endothelial cells in the dynamic blood-brain barrier-neural organoid-lab-on-a-chip (BBB-NO-LOC) model. (MACE-seq, n=4, TPM: transcript number/1 million transcripts; proteomics, n=3; mean ± SD). (A) Glycocalyx enzyme genes. (B) Glycocalyx core and enzyme proteins. (C) Drug metabolizing enzyme genes. (D) Drug metabolizing enzyme proteins. (E) Lectin genes. (F) Galectine proteins.

**Fig. S3.**
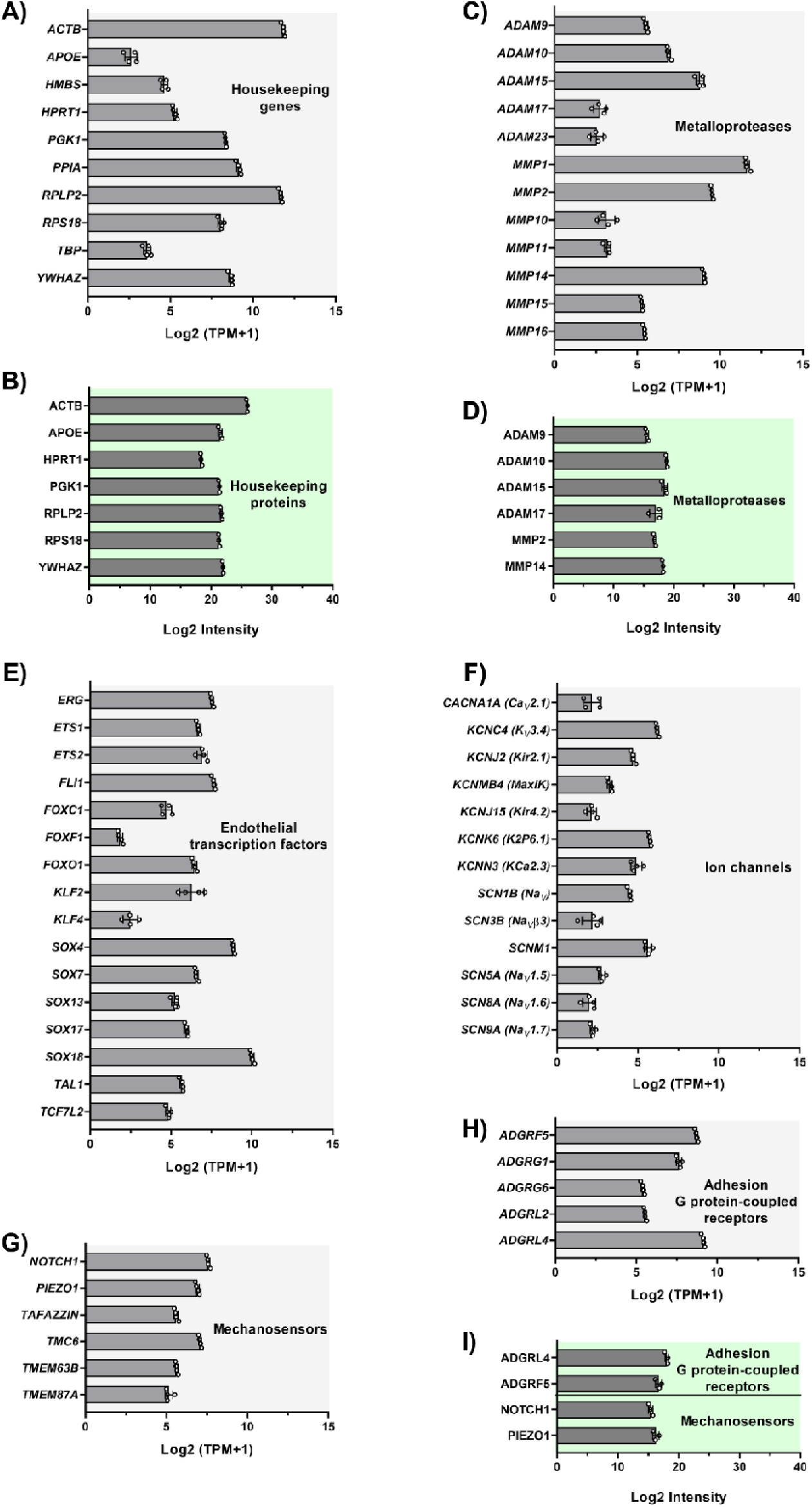
Gene- and protein expression of cultured brain endothelial cells in the dynamic blood-brain barrier-neural organoid-lab-on-a-chip (BBB-NO-LOC) model. (MACE-seq, n=4, TPM: transcript number/1 million transcripts; proteomics, n=3; mean ± SD). (A) Housekeeping genes. (B) Housekeeping proteins. (C) Metalloprotease genes. (D) Metalloprotease proteins. (E) Endothelial transcription factor genes. (F) Ion channel genes. (G) Mechanosensor genes. (H) Adhesion G-protein-coupled receptor genes. (I) Adhesion G-protein-coupled receptor and mechanosensory proteins

**Fig. S4.**
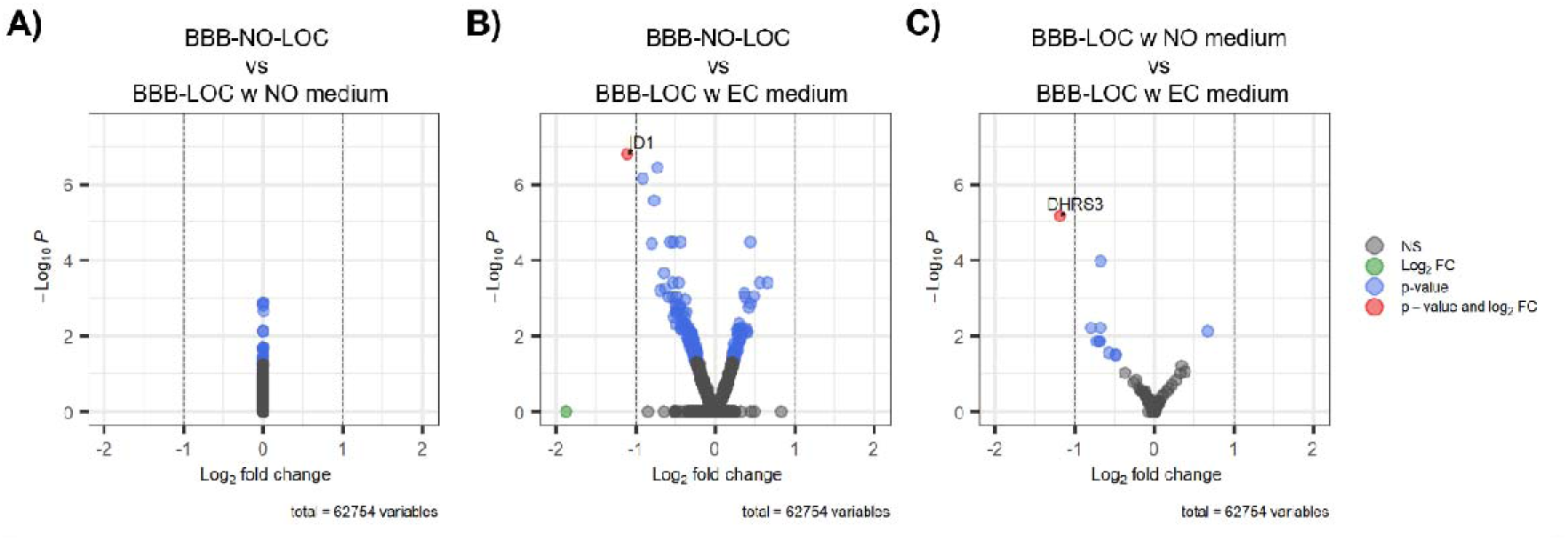
Volcano plots showing the effects of co-culture with neural organoids and culture medium composition on the transcriptome of human brain-like ECs in dynamic conditions. (A) BBB-NO-LOC model compared to BBB-LOC model without organoid, both cultured in the same culture medium (1:1 ratio of NO and EC media). (B) BBB-NO-LOC model compared to the BBB-LOC model in EC medium. (C) BBB-LOC model in 1:1 ratio of NO and EC media compared to BBB-LOC model cultured in EC medium. Dashed lines indicate thresholds of |log fold change| = 1 and *P* = 0.05. Gray, not significant; green, |log FC| > 1 only; blue, *P*<0.05 only; red, both *P*<0.05 and |log FC| > 1 (labeled: ID1 in B, DHRS3 in C). Total = 62,754 variables tested in each comparison.

**Table S2.**
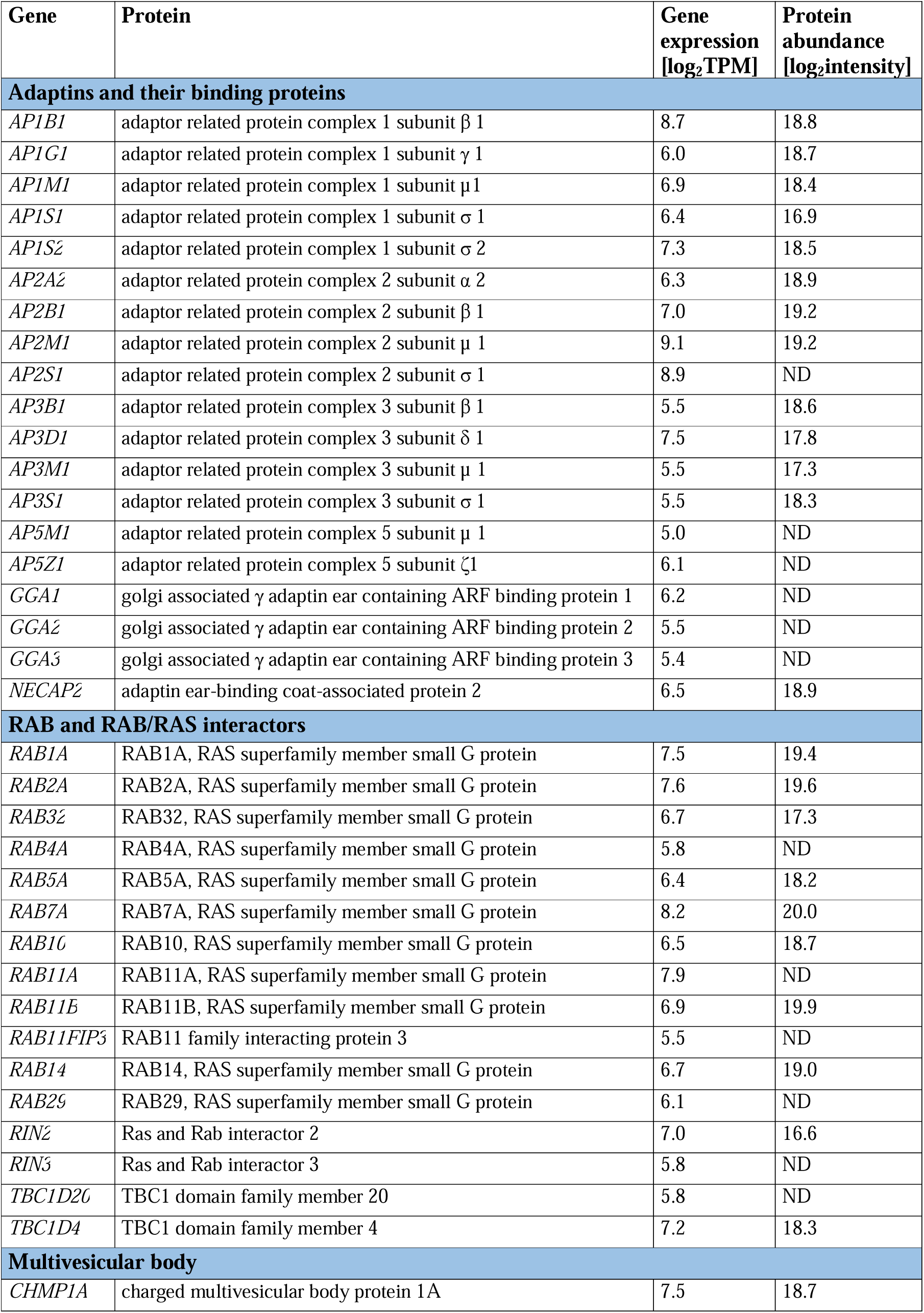
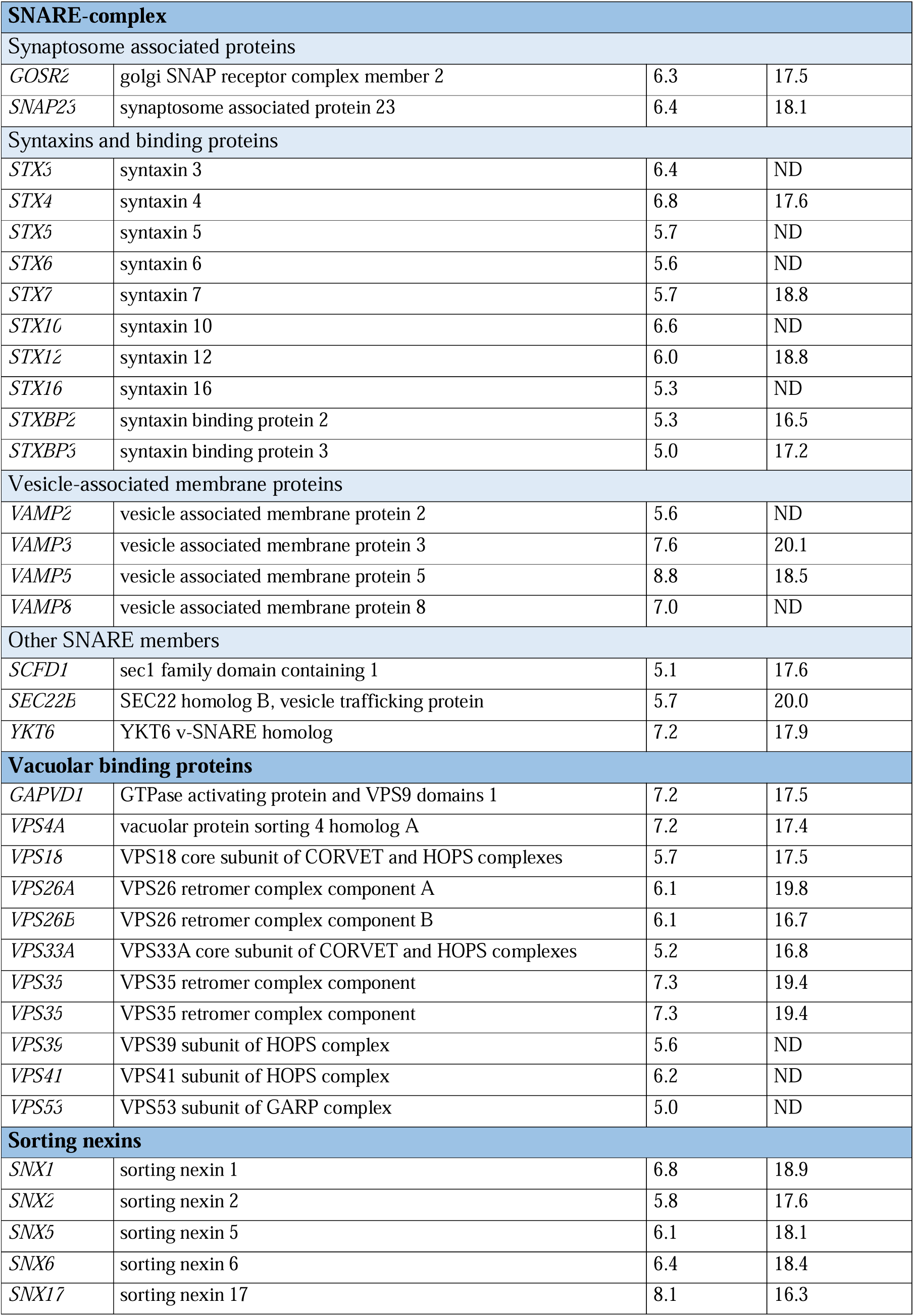
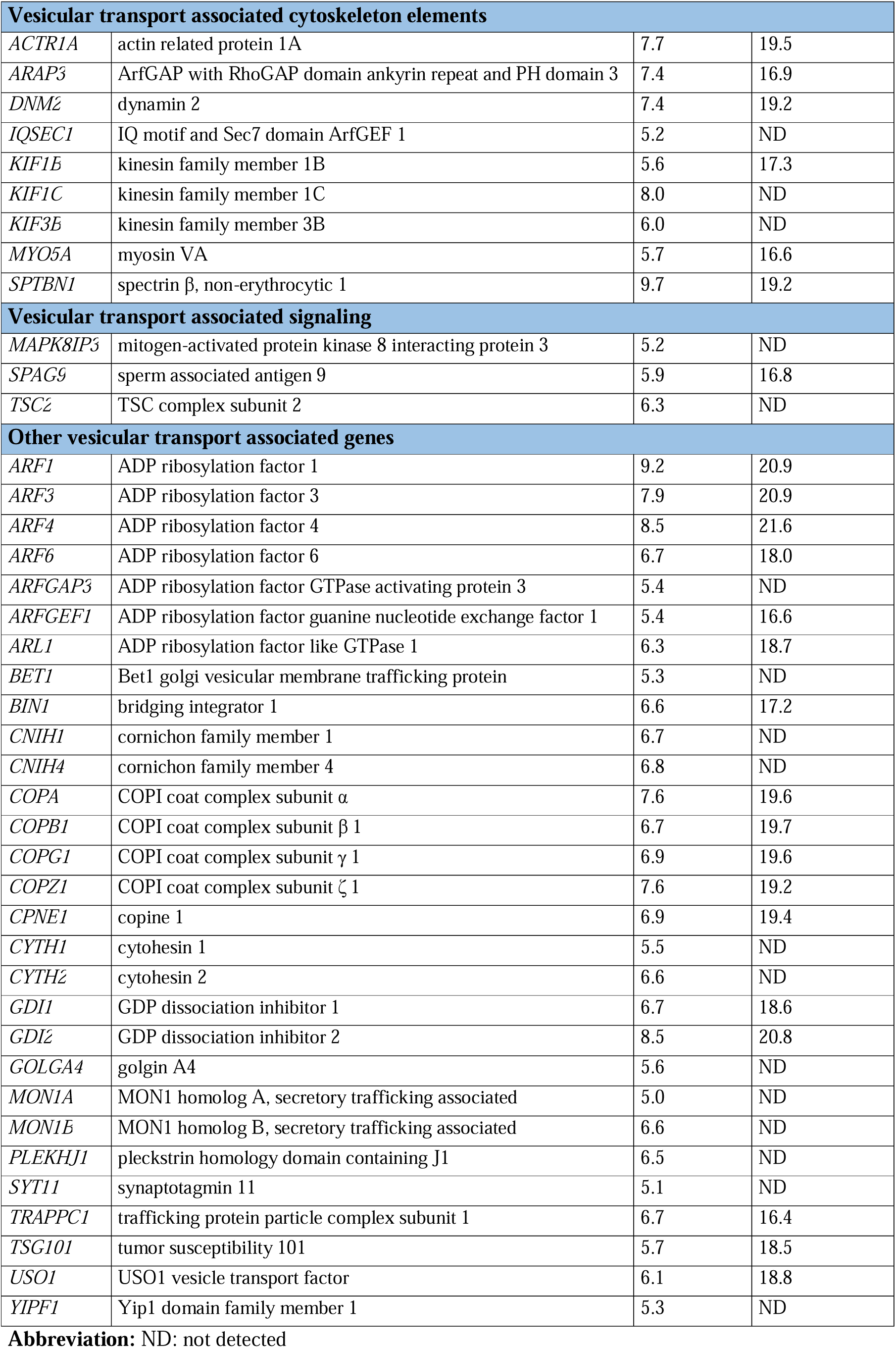
Vesicular transport associated gene- and protein expression of cultured brain endothelial cells in the dynamic blood-brain barrier-neural organoid-lab-on-a-chip model.

| Gene | Protein | Gene expression<br>[log<sub>2</sub>TPM] | Protein abundance<br>[log<sub>2</sub>intensity] |
| --- | --- | --- | --- |
| **Adaptins and their binding proteins** |  |  |  |
| *AP1B1* | adaptor related protein complex 1 subunit $\beta$ 1 | 8.7 | 18.8 |
| *AP1G1* | adaptor related protein complex 1 subunit $\gamma$ 1 | 6.0 | 18.7 |
| *AP1M1* | adaptor related protein complex 1 subunit μ1 | 6.9 | 18.4 |
| *AP1S1* | adaptor related protein complex 1 subunit $\sigma$ 1 | 6.4 | 16.9 |
| *AP1S2* | adaptor related protein complex 1 subunit $\sigma$ 2 | 7.3 | 18.5 |
| *AP2A2* | adaptor related protein complex 2 subunit $\alpha$ 2 | 6.3 | 18.9 |
| *AP2B1* | adaptor related protein complex 2 subunit $\beta$ 1 | 7.0 | 19.2 |
| *AP2M1* | adaptor related protein complex 2 subunit $\mu$ 1 | 9.1 | 19.2 |
| *AP2S1* | adaptor related protein complex 2 subunit $\sigma$ 1 | 8.9 | ND |
| *AP3B1* | adaptor related protein complex 3 subunit $\beta$ 1 | 5.5 | 18.6 |
| *AP3D1* | adaptor related protein complex 3 subunit $\delta$ 1 | 7.5 | 17.8 |
| *AP3M1* | adaptor related protein complex 3 subunit $\mu$ 1 | 5.5 | 17.3 |
| *AP3S1* | adaptor related protein complex 3 subunit $\sigma$ 1 | 5.5 | 18.3 |
| *AP5M1* | adaptor related protein complex 5 subunit $\mu$ 1 | 5.0 | ND |
| *AP5Z1* | adaptor related protein complex 5 subunit ζ1 | 6.1 | ND |
| *GGA1* | golgi associated $\gamma$ adaptin ear containing ARF binding protein 1 | 6.2 | ND |
| *GGA2* | golgi associated $\gamma$ adaptin ear containing ARF binding protein 2 | 5.5 | ND |
| *GGA3* | golgi associated $\gamma$ adaptin ear containing ARF binding protein 3 | 5.4 | ND |
| *NECAP2* | adaptin ear-binding coat-associated protein 2 | 6.5 | 18.9 |
| **RAB and RAB/RAS interactors** |  |  |  |
| *RAB1A* | RAB1A, RAS superfamily member small G protein | 7.5 | 19.4 |
| *RAB2A* | RAB2A, RAS superfamily member small G protein | 7.6 | 19.6 |
| *RAB32* | RAB32, RAS superfamily member small G protein | 6.7 | 17.3 |
| *RAB4A* | RAB4A, RAS superfamily member small G protein | 5.8 | ND |
| *RAB5A* | RAB5A, RAS superfamily member small G protein | 6.4 | 18.2 |
| *RAB7A* | RAB7A, RAS superfamily member small G protein | 8.2 | 20.0 |
| *RAB10* | RAB10, RAS superfamily member small G protein | 6.5 | 18.7 |
| *RAB11A* | RAB11A, RAS superfamily member small G protein | 7.9 | ND |
| *RAB11B* | RAB11B, RAS superfamily member small G protein | 6.9 | 19.9 |
| *RAB11FIP3* | RAB11 family interacting protein 3 | 5.5 | ND |
| *RAB14* | RAB14, RAS superfamily member small G protein | 6.7 | 19.0 |
| *RAB29* | RAB29, RAS superfamily member small G protein | 6.1 | ND |
| *RIN2* | Ras and Rab interactor 2 | 7.0 | 16.6 |
| *RIN3* | Ras and Rab interactor 3 | 5.8 | ND |
| *TBC1D20* | TBC1 domain family member 20 | 5.8 | ND |
| *TBC1D4* | TBC1 domain family member 4 | 7.2 | 18.3 |
| **Multivesicular body** |  |  |  |
| *CHMP1A* | charged multivesicular body protein 1A | 7.5 | 18.7 |

| **SNARE-complex** |  |  |  |
| --- | --- | --- | --- |
| Synaptosome associated proteins |  |  |  |
| *GOSR2* | golgi SNAP receptor complex member 2 | 6.3 | 17.5 |
| *SNAP23* | synaptosome associated protein 23 | 6.4 | 18.1 |
| Syntaxins and binding proteins |  |  |  |
| *STX3* | syntaxin 3 | 6.4 | ND |
| *STX4* | syntaxin 4 | 6.8 | 17.6 |
| *STX5* | syntaxin 5 | 5.7 | ND |
| *STX6* | syntaxin 6 | 5.6 | ND |
| *STX7* | syntaxin 7 | 5.7 | 18.8 |
| *STX10* | syntaxin 10 | 6.6 | ND |
| *STX12* | syntaxin 12 | 6.0 | 18.8 |
| *STX16* | syntaxin 16 | 5.3 | ND |
| *STXBP2* | syntaxin binding protein 2 | 5.3 | 16.5 |
| *STXBP3* | syntaxin binding protein 3 | 5.0 | 17.2 |
| Vesicle-associated membrane proteins |  |  |  |
| *VAMP2* | vesicle associated membrane protein 2 | 5.6 | ND |
| *VAMP3* | vesicle associated membrane protein 3 | 7.6 | 20.1 |
| *VAMP5* | vesicle associated membrane protein 5 | 8.8 | 18.5 |
| *VAMP8* | vesicle associated membrane protein 8 | 7.0 | ND |
| Other SNARE members |  |  |  |
| *SCFD1* | sec1 family domain containing 1 | 5.1 | 17.6 |
| *SEC22B* | SEC22 homolog B, vesicle trafficking protein | 5.7 | 20.0 |
| *YKT6* | YKT6 v-SNARE homolog | 7.2 | 17.9 |
| **Vacuolar binding proteins** |  |  |  |
| *GAPVD1* | GTPase activating protein and VPS9 domains 1 | 7.2 | 17.5 |
| *VPS4A* | vacuolar protein sorting 4 homolog A | 7.2 | 17.4 |
| *VPS18* | VPS18 core subunit of CORVET and HOPS complexes | 5.7 | 17.5 |
| *VPS26A* | VPS26 retromer complex component A | 6.1 | 19.8 |
| *VPS26B* | VPS26 retromer complex component B | 6.1 | 16.7 |
| *VPS33A* | VPS33A core subunit of CORVET and HOPS complexes | 5.2 | 16.8 |
| *VPS35* | VPS35 retromer complex component | 7.3 | 19.4 |
| *VPS35* | VPS35 retromer complex component | 7.3 | 19.4 |
| *VPS39* | VPS39 subunit of HOPS complex | 5.6 | ND |
| *VPS41* | VPS41 subunit of HOPS complex | 6.2 | ND |
| *VPS53* | VPS53 subunit of GARP complex | 5.0 | ND |
| **Sorting nexins** |  |  |  |
| *SNX1* | sorting nexin 1 | 6.8 | 18.9 |
| *SNX2* | sorting nexin 2 | 5.8 | 17.6 |
| *SNX5* | sorting nexin 5 | 6.1 | 18.1 |
| *SNX6* | sorting nexin 6 | 6.4 | 18.4 |
| *SNX17* | sorting nexin 17 | 8.1 | 16.3 |

| Vesicular transport associated cytoskeleton elements |  |  |  |
| --- | --- | --- | --- |
| *ACTR1A* | actin related protein 1A | 7.7 | 19.5 |
| *ARAP3* | ArfGAP with RhoGAP domain ankyrin repeat and PH domain 3 | 7.4 | 16.9 |
| *DNM2* | dynamain 2 | 7.4 | 19.2 |
| *IQSEC1* | IQ motif and Sec7 domain ArfGEF 1 | 5.2 | ND |
| *KIF1B* | kinesin family member 1B | 5.6 | 17.3 |
| *KIF1C* | kinesin family member 1C | 8.0 | ND |
| *KIF3B* | kinesin family member 3B | 6.0 | ND |
| *MYO5A* | myosin VA | 5.7 | 16.6 |
| *SPTBN1* | spectrin $\beta$, non-erythrocytic 1 | 9.7 | 19.2 |
| Vesicular transport associated signaling |  |  |  |
| *MAPK8IP3* | mitogen-activated protein kinase 8 interacting protein 3 | 5.2 | ND |
| *SPAG9* | sperm associated antigen 9 | 5.9 | 16.8 |
| *TSC2* | TSC complex subunit 2 | 6.3 | ND |
| Other vesicular transport associated genes |  |  |  |
| *ARF1* | ADP ribosylation factor 1 | 9.2 | 20.9 |
| *ARF3* | ADP ribosylation factor 3 | 7.9 | 20.9 |
| *ARF4* | ADP ribosylation factor 4 | 8.5 | 21.6 |
| *ARF6* | ADP ribosylation factor 6 | 6.7 | 18.0 |
| *ARFGAP3* | ADP ribosylation factor GTPase activating protein 3 | 5.4 | ND |
| *ARFGEF1* | ADP ribosylation factor guanine nucleotide exchange factor 1 | 5.4 | 16.6 |
| *ARL1* | ADP ribosylation factor like GTPase 1 | 6.3 | 18.7 |
| *BET1* | Bet1 golgi vesicular membrane trafficking protein | 5.3 | ND |
| *BIN1* | bridging integrator 1 | 6.6 | 17.2 |
| *CNIH1* | cornichon family member 1 | 6.7 | ND |
| *CNIH4* | cornichon family member 4 | 6.8 | ND |
| *COPA* | COPI coat complex subunit $\alpha$ | 7.6 | 19.6 |
| *COPB1* | COPI coat complex subunit $\beta$ 1 | 6.7 | 19.7 |
| *COPG1* | COPI coat complex subunit $\gamma$ 1 | 6.9 | 19.6 |
| *COPZ1* | COPI coat complex subunit $\zeta$ 1 | 7.6 | 19.2 |
| *CPNE1* | copine 1 | 6.9 | 19.4 |
| *CYTH1* | cytohesin 1 | 5.5 | ND |
| *CYTH2* | cytohesin 2 | 6.6 | ND |
| *GDI1* | GDP dissociation inhibitor 1 | 6.7 | 18.6 |
| *GDI2* | GDP dissociation inhibitor 2 | 8.5 | 20.8 |
| *GOLGA4* | golgin A4 | 5.6 | ND |
| *MON1A* | MON1 homolog A, secretory trafficking associated | 5.0 | ND |
| *MON1B* | MON1 homolog B, secretory trafficking associated | 6.6 | ND |
| *PLEKHJ1* | pleckstrin homology domain containing J1 | 6.5 | ND |
| *SYT11* | synaptotagmin 11 | 5.1 | ND |
| *TRAPPC1* | trafficking protein particle complex subunit 1 | 6.7 | 16.4 |
| *TSG101* | tumor susceptibility 101 | 5.7 | 18.5 |
| *USO1* | USO1 vesicle transport factor | 6.1 | 18.8 |
| *YIPF1* | Yip1 domain family member 1 | 5.3 | ND |

**Abbreviation:** ND: not detected

**Table S3.** Physico-chemical properties of non-targeted (N) and targeted (N-A-GSH) niosomes containing Texas red-labeled bovine serum albumin as cargo.

|  | Diameter<br>(nm) | PDI | Zeta potential<br>(mV) | Encapsulation<br>efficiency (%) |
| --- | --- | --- | --- | --- |
| **N** | 92.61 ± 2.07 | 0.25 ± 0.01 | -8.53 ± 0.35 | 22.88%<br>111.45 µg/ml |
| **N-A-GSH** | 81.02 ± 1.67 | 0.25 ± 0.01 | -9.05 ± 0.07 | 20.39%<br>89.68 µg/ml |

**Abbreviation:** PDI: polydispersity index.

**Fig. S5.**
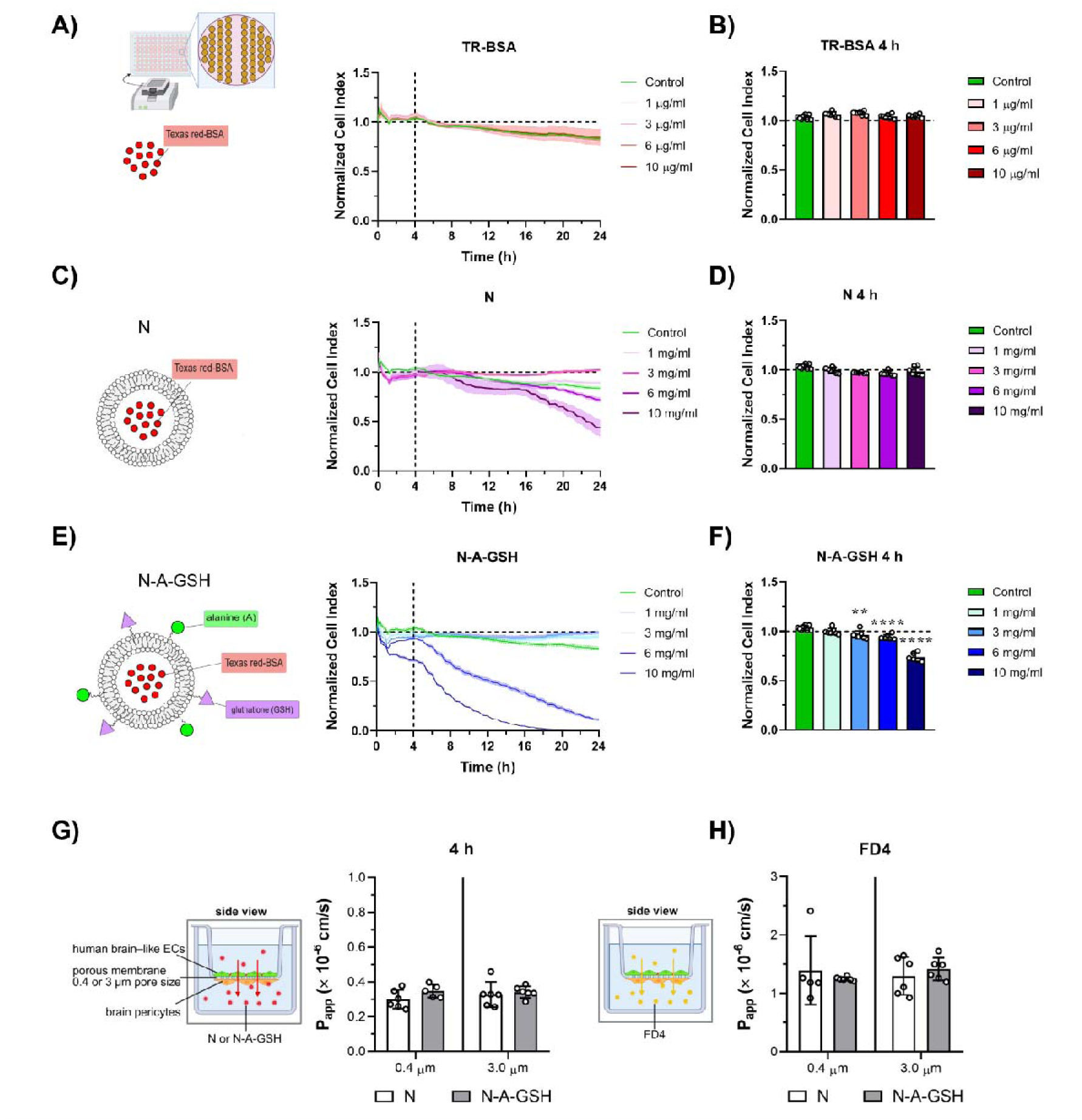
Cell response of human brain-like endothelial cells to non-targeted (N) and alanine-glutathione targeted (N-A-GSH) niosomes containing Texas-red labeled bovine serum albumin (TR-BSA) as cargo. (A) Real-time impedance measurement of human brain-like EC monolayers incubated with TR-BSA for 24h. (B) Impedance (cell index) of human brain-like EC monolayers incubated with TR-BSA at the 4-hour time-point. (C) Real-time impedance measurement of human brain-like EC monolayers incubated with TR-BSA loaded non-targeted niosomes (N) for 24h. (D) Impedance of human brain-like EC monolayers incubated with N at the 4-hour time-point. (E) Real-time impedance measurement of human brain-like EC monolayers incubated with N-A-GSH for 24h. (F) Impedance of human brain-like EC monolayers incubated with N-A-GSH at the 4-hour time-point. (A-F) Mean ± SD, one-way ANOVA, Bonferroni post-test, n = 4-8, ** P<0.01; **** P<0.0001. (G) Apparent permeability coefficients (P_app_) of N and N-A-GSH across the contact co-culture human BBB model on inserts with 0.4 and 3 µm pore size membranes for 4h. (H) P_app_ of fluorescein isothiocyanate labeled 4 kDa dextran (FD4) on the human BBB model assembled on inserts with 0.4 and 3 µm pore size membranes following the 4-hour nanoparticle assay. (G-H) Mean ± SD, two-way ANOVA, Bonferroni post-test, n = 5-6, no statistical significance.

**Fig. S6.**
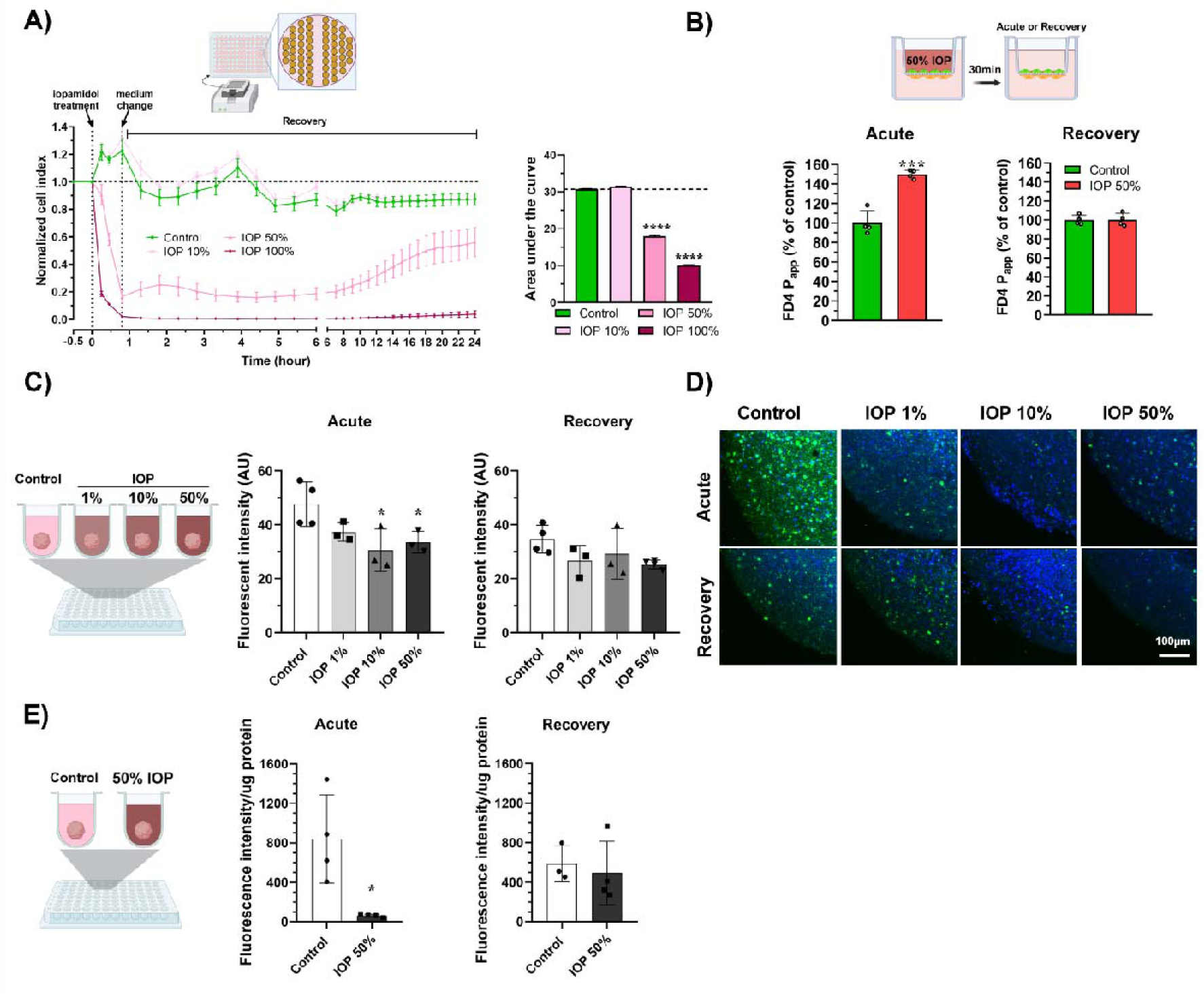
Effect of iodinated hyperosmolar contrast agent iopamidol (IOP) on cultured brain-like endothelial cells (ECs) and neural organoids. (A) Real-time impedance measurement of human brain-like EC monolayers after 30 min IOP treatment and 24 hours recovery with normal EC medium. Normalized cell index and area under the curve. One-way ANOVA, Bonferroni post-test, mean ± SD, n = 9, **** P<0.001. (B) Fluorescein isothiocyanate 4 kDa (FD4) permeability on the static BBB model after 30 min acute IOP treatment and after 24h recovery with EC medium. Unpaired t-test, mean ± SD, n = 4, *** P<0.001. (C) Effect of direct IOP treatment on the metabolic activity of neural organoids. After 30 min acute treatment and 24 h recovery calcein AM assay was performed. Mean ± SD, one-way ANOVA, Bonferroni post-test, n = 3-4, * P<0.05. (D) Representative images of neural organoids after the calcein AM assay. Green: calcein, blue: cell nuclei (H33342). Scale bar: 100 µm (E) Effect of direct IOP treatment on the esterase activity of neural organoids. Unpaired t-test, mean ± SD, n = 3-4, * P<0.05.

